# Genome Sequencing Unveils a New Regulatory Landscape of Platelet Reactivity

**DOI:** 10.1101/621565

**Authors:** Ali R. Keramati, Ming-Huei Chen, Benjamin A.T. Rodriguez, Lisa R. Yanek, Brady J. Gaynor, Kathleen Ryan, Jennifer A. Brody, NHLBI Trans-Omics for Precision (TOPMed) Consortium, NHLBI TOPMed Hematology and Hemostasis Working Group, Kai Kammers, Kanika Kanchan, Kruthika Iyer, Madeline H. Kowalski, Achilleas N. Pitsillides, L. Adrienne Cupples, Alan R. Shuldiner, Jeffrey R. O’Connell, Braxton D. Mitchell, Nauder Faraday, Margaret A. Taub, Lewis C. Becker, Joshua P. Lewis, Rasika A. Mathias, Andrew D. Johnson

## Abstract

Exaggerated platelet aggregation at the site of vascular injury is the underlying pathophysiology of thrombotic diseases. Here, we conduct the largest whole genome sequencing (WGS) effort to uncover the genetic determinants of platelet aggregation. Leveraging 3,855 NHLBI Trans-Omics for Precision Medicine (TOPMed) individuals deeply phenotyped for platelet aggregation, we identify 18 loci using single-variant approaches. This includes the novel RGS18 locus encoding a myeloerythroid lineage-specific regulator of G-protein signaling that co-localizes with eQTL signatures for RGS18 expression in platelets. A gene-based approach focusing on deleterious coding variants identifies the SVEP1 gene, previously shown to be associated with coronary artery disease, as a novel determinant of platelet aggregation. Finally, in an integrative approach leveraging epigenetic data on megakaryocytes, we find strong association between rare variants mapping to a super enhancer region for PEAR1. This is a novel finding implicating the importance of rare variants with regulatory potential in a previously documented GWAS-identified locus.

---

Platelet aggregation at the site of atherosclerotic vascular injury is the underlying pathophysiology of myocardial infarction and stroke ^1^. Identification of the factors that contribute to increased platelet reactivity is critical to understanding the biology of arterial thrombus formation and in the development of novel treatment strategies for patients at risk for pathologic vascular thrombosis. Here, we conduct the largest whole genome sequencing (WGS) effort to uncover the genetic determinants of platelet aggregation by leveraging 3,855 NHLBI Trans-Omics for Precision Medicine (TOPMed) participants deeply phenotyped for platelet aggregation. Using single-variant approaches, we identify 16 independent loci including the novel *RGS18* locus, which encodes a myeloerythroid lineage-specific regulator of G-protein signaling that co-localizes with expression quantitative trait loci (eQTL) signatures for *RGS18* expression in platelets. Additionally, a gene-based approach focusing on deleterious coding variants identifies the *SVEP1* gene, a known contributor of coronary artery disease risk, as a novel determinant of platelet aggregation. Finally, in an integrative approach leveraging epigenetic data on megakaryocytes (MK), we observe strong association between rare variants mapping to a super-enhancer region for *PEAR1* and platelet aggregation. Together, these results unveil novel loci contributing to platelet reactivity, provide insights regarding the mechanism(s) by which genetic variation may influence cardiovascular disease risk, and underscore the importance of utilizing rare variant and regulatory approaches to better understand loci contributing to complex phenotypes.

Previous studies have shown that platelet aggregation in response to agonists is highly heritable ^2–6^ and prior genome-wide association studies (GWAS) have identified 7 common variants for platelet aggregation in response to different agonists.^7–9^ Using a WGS approach, we aimed to i) refine previously identified GWAS loci; ii) identify novel loci; iii) examine the burden of deleterious coding variants; and iv) evaluate rare non-coding variants within MK-specific super-enhancer regions for 19 harmonized phenotypic measures of platelet aggregation in response to different doses of adenosine diphosphate (ADP), epinephrine, and collagen (***Supplementary Table 1***).

Genome-wide single variant tests for association were performed on ∼28 million variants in 3,125 European Americans (EA) and 730 African Americans (AA) (***Supplementary Table 2***) from the Framingham Heart Study (FHS), Older Order Amish Study (OOA), and the Genetic Study of Atherosclerosis Risk (GeneSTAR). There was no evidence of test statistic inflation using a mega-analysis approach combining all samples (***Supplementary Figure 1***). We identified 101 variants associated with platelet aggregation in response to ADP, epinephrine, or collagen (P-value<5×10^−8^, ***Figure 1***). Using iterative conditional analyses, genome-wide significant variants were refined down to 16 independent loci (***Table 1***). With the exception of two variants (rs12041331 and chr17:21960955) all of these loci were uniquely associated with platelet aggregation in response to one agonist, and most of the identified loci were novel (***Table 1, Supplementary Figure 2***). Replication of discovery findings was performed in up to 2,009 additional samples imputed to the TOPMed reference panel that were not in the discovery analyses (***Supplementary Table 3a***), and extended to the Caerphilly Prospective Study [CaPS] (N=1,183) for ADP and collagen-induced platelet aggregation phenotypes^8,10^ (***Supplementary Table 4***).

**Table 1:** Sixteen loci identified through single variant approaches for genome-wide association for platelet aggregation in response to epinephrine, ADP and collagen in 3,855 TOPMed participants. P-values presented are a summary across all individual phenotypes for the single agonist (i.e. the minimum P-value for the SNV from 8, 7 and 4 individual phenotypes for epinephrine, ADP, and collagen, respectively) (see **Supplementary Table 1**).

| Known<br>vs Novel | hg38 | rsID | ref/alt | Nearest Gene | MAF | ADP |  | Epinephrine |  | Collagen |  |
| --- | --- | --- | --- | --- | --- | --- | --- | --- | --- | --- | --- |
|  |  |  |  |  |  | P-value | Beta | P-value | Beta | P-value | Beta |
| Novel | Chr1:192194880 | rs1175170 | G/C | RGS18,RGS21 | 0.442 | 7.86E-06 | 0.123 | 1.96E-09 | 0.154 | 2.37E-02 | -0.056 |
| Known | Chr1:156899922 | rs12041331 | G/A | PEAR1 | 0.148 | 7.61E-17 | -0.329 | 2.30E-18 | -0.357 | 7.57E-17 | 0.317 |
| Novel | Chr1:20567949 | rs12137738 | A/T | FAM43B,CDA | 0.077 | 0.0002 | 0.219 | 1.04E-08 | 0.305 | 0.017 | -0.107 |
| Novel | Chr1:67128641 | rs142001088 | C/T | C1orf141 | 0.018 | 0.0281 | 0.203 | 2.51E-03 | 0.275 | 9.25E-09 | -0.502 |
| Novel | Chr5:19109993 | rs112157462 | T/C | LINC02223,CDH18 | 0.022 | 1.64E-02 | -0.281 | 5.90E-03 | -0.229 | 1.18E-08 | 0.457 |
| Novel | Chr6:121921871 | rs58250884 | A/G | GJA1,HSF2 | 0.087 | 1.60E-03 | -0.153 | 2.22E-08 | -0.272 | 2.34E-02 | 0.105 |
| Novel | Chr9:28873884 | rs185159562 | T/A | LINGO2 | 0.005 | 3.87E-08 | -0.987 | 5.10E-03 | -0.987 | 1.45E-01 | 0.243 |
| Novel | Chr10:75490891 | rs138028657 | A/G | LRMDA | 0.006 | 0.6443 | 0.072 | 0.6144 | -0.1017 | 1.52E-08 | -0.858 |
| Known | Chr10:111139289 | rs7097060 | T/A | ADRA2A,GPAM | 0.137 | 5.45E-01 | 0.028 | 6.68E-12 | 0.251 | 6.73E-01 | -0.014 |
| Novel | Chr11:92185065 | rs183146849 | A/T | DISC1FP1,FAT3 | 0.012 | 3.10E-08 | -0.702 | 1.33E-02 | -0.375 | 4.38E-01 | -0.080 |
| Novel | Chr12:132589485 | rs140148392 | G/A | FBRSL1,LRCOL1 | 0.009 | 1.51E-01 | 0.253 | 8.00E-03 | 0.137 | 2.88E-08 | 0.668 |
| Novel | Chr13:96912429 | rs61974290 | A/G | HS6ST3,LINC00359 | 0.057 | 9.57E-02 | 0.088 | 3.36E-09 | -0.399 | 9.40E-03 | 0.133 |
| Novel | Chr17:21960955 | . | A/T | KCNJ18,UBBP4 | 0.276 | 2.87E-08 | 0.167 | 7.73E-19 | 0.259 | 1.80E-03 | -0.089 |
| Novel | Chr17:16451482 | rs575524466 | G/A | LRRC75A-AS1 | 0.003 | 1.90E-01 | -0.315 | 2.60E-03 | -0.462 | 3.12E-08 | 1.168 |
| Novel | Chr18:29059923 | rs138845468 | TAAATA/T | CDH2,MIR302F | 0.082 | 6.90E-02 | 0.087 | 1.21E-01 | 0.086 | 3.22E-08 | -0.250 |
| Novel | Chr20:50142397 | rs542707094 | CTG/C | TMEM189,TMEM189-UBE2V1 | 0.003 | 3.16E-02 | -0.485 | 3.05E-02 | -0.486 | 3.52E-08 | 1.193 |

**Figure 1:**
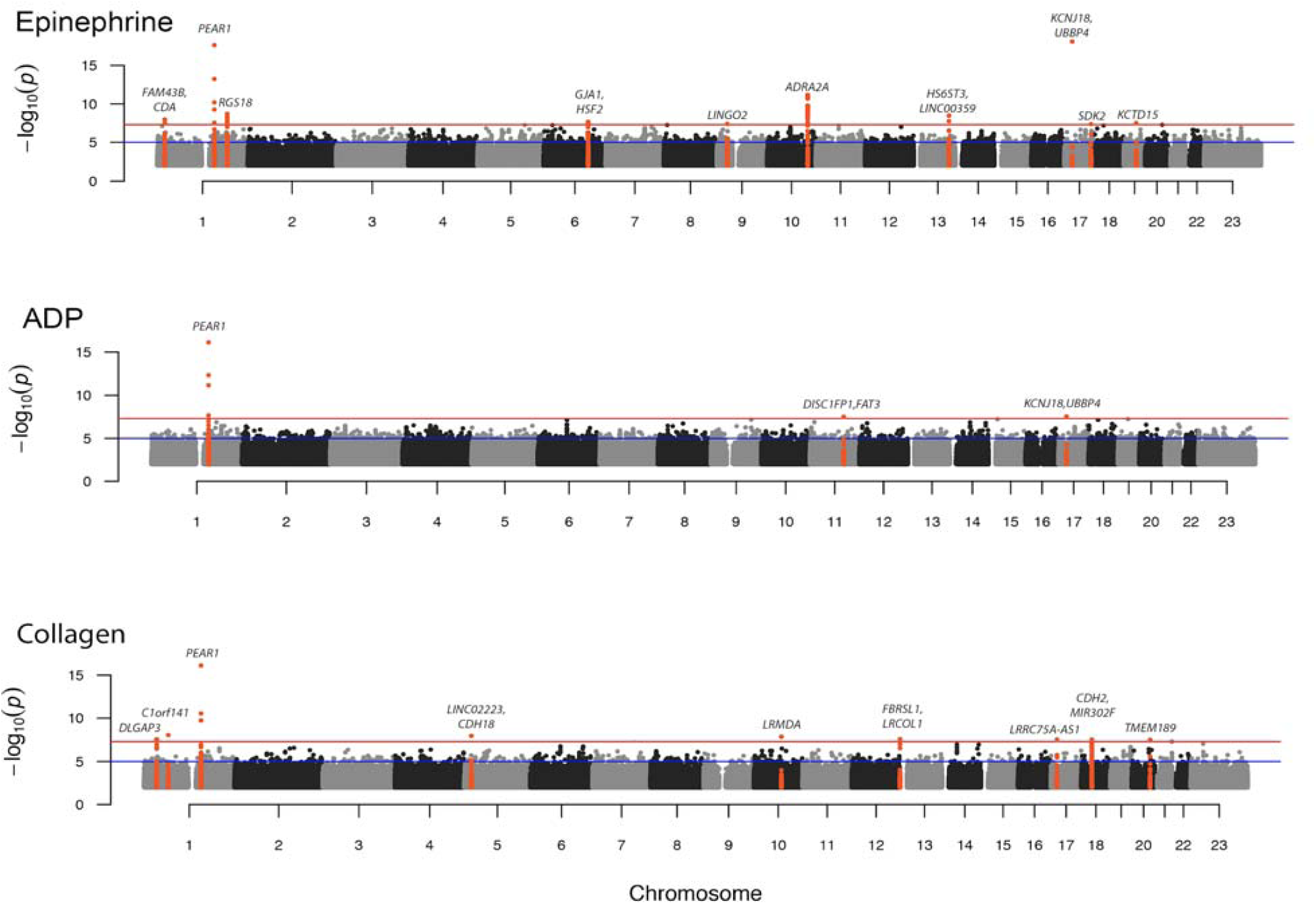
Genome-wide association study results for platelet aggregation in response to epinephrine, ADP and collagen in 3,855 TOPMed participants. P-values presented are a summary across all individual phenotypes for the single agonist (i.e. the minimum P-value for the variants from 8, 7 and 4 individual phenotypes for epinephrine, ADP, and collagen, respectively) (**Supplementary Table 1**). Loci passing genome wide significance (5×10^−8^) are marked by red dots. Locus names represent the nearest (for novel) or previously annotated (for prior) gene. The blue horizontal line indicates a P-value threshold of 1×10^−5^ corresponding to suggestive significance threshold and the red line indicates a P-value threshold of 5×10^−8^, corresponding to genome-wide significance.

**Figure 2:**
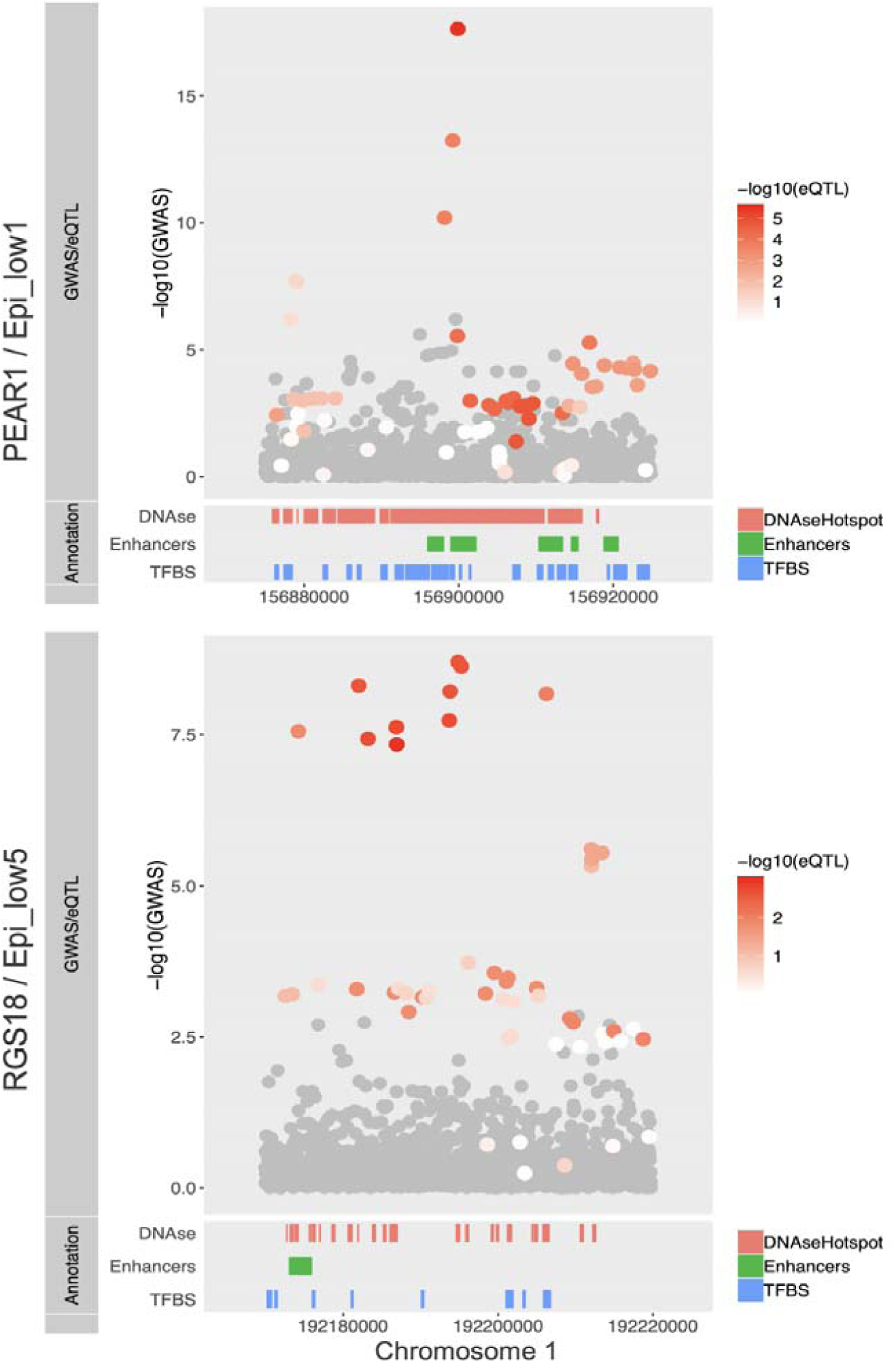
Co-localization of WGS association signal and platelet eQTL signatures. Top panel shows co-localization between *PEAR1* platelet eQTL signatures and WGS association evidence for platelet aggregation in response to Epi_low1 (see **Supplementary Table 1**) on Chr1:156899922 and bottom panel is between *RGS18* platelet eQTL signatures with WGS association evidence for platelet aggregation in response to Epi_low 5 (see Supplementary **Table 1**) on Chr1:156899922. Region of co-localization is zoomed to peak SNV +/- 25kb, strength of the eQTL evidence for the gene is shown as a heat scale in red, SNVs that were not included in eQTL analysis are shown in gray. Annotations of regulatory elements include ENCODE-aggregated cell model data for both open chromatin (DNAse, Red) as well as transcription factor binding clusters (TFBS, Blue) and cell type-specific megakaryocyte distal enhancer elements (Enhancers, Green).

Given that all of the 16 loci identified using single-variant approaches are located in non-coding regions of the genome, we tested for co-localization between these regions and eQTL data available through RNA sequencing of platelets in 180 European Americans from GeneSTAR (***Supplementary Table 5***). We found that sentinel variants in the PEAR1 and RGS18 loci were eQTLs for *PEAR1* and *RGS18*, respectively. Furthermore, co-localization signatures for *PEAR1* and *RGS18* were identified using Bayesian approaches (***Figure 2***). Previous work demonstrated a single, common (∼14% MAF) intronic peak variant in *PEAR1* (rs12041331) is associated with platelet phenotypes using GWAS, regional sequencing, and exonic approaches ^8,11–13^. The minor allele of rs12041331 is linked to decreased *PEAR1* platelet protein levels ^13,14^, potentially through alteration of a methylation site in MKs^15^. In addition, the role of this gene in platelet signaling is supported by mechanistic studies ^16,17^. Our co-localization results with platelet eQTLs are consistent with a model that a single, common causal variant explains the platelet reactivity signal. While results of previous association studies on *PEAR1* variants and coronary disease are mixed ^18–21^, UK BioBank ^22^ data supports an association (***Supplementary Table 6***).

In the novel RGS18 region, several variants were replicated using independent samples imputed with the TOPMed panel (***Supplementary Table 3***), and independent evidence was also found for ADP and collagen aggregation phenotypes in the CaPS study (***Supplementary Table 4***). Overlaying the associated variants with platelet eQTLs and megakaryocytic epigenome features, there are several potential candidate variants (***Supplementary Table 7***). Consistent with our human results, independent *Rgs18*^-/-^ mouse studies suggest Rgs18 inhibits pre-agonist stimulated platelet reactivity, with knockouts exhibiting exaggerated platelet reactivity to multiple agonist pathways, decreased bleeding times, and increased arterial occlusion ^23,24^. This is attributed to a loss of inhibition of multiple G-protein coupled receptor signaling pathways in platelets ^25^. Notably, variants in *RGS18* identified in our study are also associated with altered risk for cerebral infarction and subarachnoid hemorrhage in the UK BioBank (***Supplementary Table 6***).

SKAT^26^ gene-based tests using a MAF threshold of 0.05 were conducted for deleterious variants mapping to 17,774 protein-coding genes (***Supplementary Table 8, Supplementary Figure 3***) with significant findings after Bonferroni correction for *SVEP1* (ADP-induced platelet aggregation, P-value=2.6×10^−6^), *BCO1* (epinephrine-induced platelet aggregation, P=8.9×10^−7^), *NELFA* (collagen-induced platelet aggregation, P=1.7×10^−6^) and *IDH3A* (collagen-induced platelet aggregation, P-value=2.6×10^−6^). Using a *leave-one-out* analysis, we observed that these associations were driven mainly by single or limited sets of rare variants (***Supplementary Figure 4, Supplementary Table 9***). For example, the *SVEP1* association with ADP-induced platelet aggregation was solely driven by a nonsynonymous variant (Gly229Arg) in the second exon (rs61751937, MAF 0.028, P-value=5.8×10^−6^). This variant alters a highly conserved residue located in the protein’s VWFa domain (***Figure 3a-c***). The finding replicated in the WGS imputed cohorts (P-value =0.004) and CaPS (P-value=0.008) demonstrating an association with increased ADP-induced platelet reactivity. Interestingly, a different variant in *SVEP1*, rs111245230, is associated with hypertension and increased cardiovascular risk in an exome study ^27^. This variant is in linkage equilibrium (r^2^=0.0002; D’=0.01) with our peak variant and not associated with ADP-induced platelet aggregation (P=0.06). Both variants are modestly associated with CVD outcomes in the UK BioBank (***Supplementary Table 10***). Collectively, these results suggest that *SVEP1* may have previously unappreciated and multifactorial roles in contributing to CVD. Homozygous *Svep1*^*-/-*^ mice die from edema, and heterozygous mice, as well as zebrafish, experience arterial and lymphatic vessel malformations ^28–30^. Consistent with previous investigations ^31^, RNAseq data in a subset of GeneSTAR participants do not indicate expression of *SVEP1* in platelets. The protein is strongly conserved (***Figure 3c***) and could potentially affect platelet function and CVD through several mechanisms including cell-cell adhesion, cell differentiation, and functions in bone marrow niches ^32^.

**Figure 3a-c:**
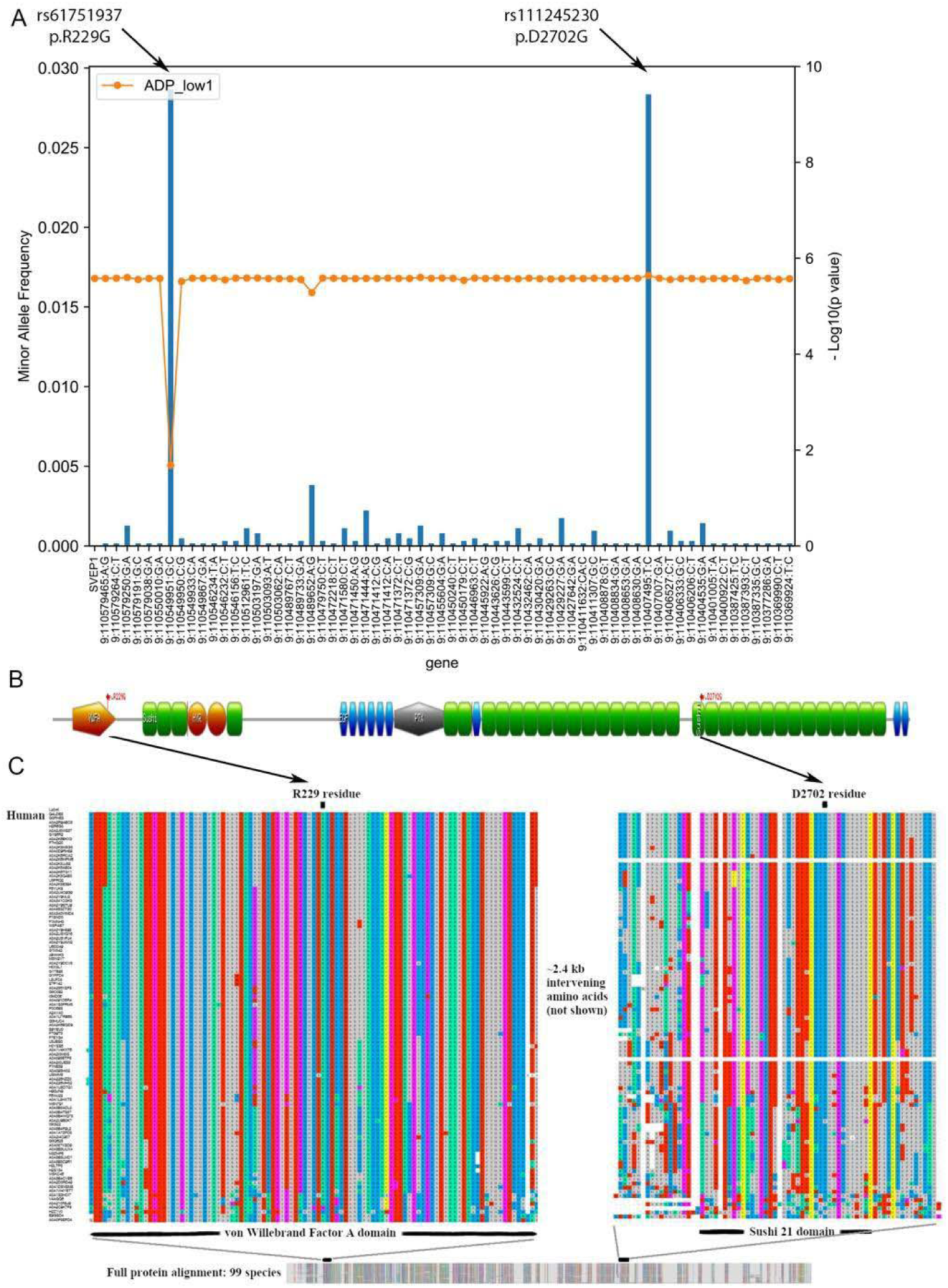
Association of aggregated rare deleterious coding variants in *SVEP1* and ADP-induced platelet aggregation. **3a**. Using a *leave-one-out* approach, we identified a rare coding variant (rs61751937) that explains most of the association. First orange dot represents the overall gene based test including 64 variants. Subsequent orange dots represent the -Log10(P-value) of the gene based test when the specific labeled variant was left out, and blue bars represent minor allele frequency (MAF) of specific variant being left out. **3b**. Schematic protein structure of SVEP1. rs61751937 substitutes glycine for arginine at position 229. Another variant in *SVEP1* has been associated with coronary artery disease which substitutes glycine for aspartic acid at position 2702. **3c**. Using UniProt, a total of 98 orthologs were identified for the largest human SVEP1 protein isoform and aligned. Alignments were visualized in MAFFT (v.7, https://mafft.cbrc.jp/alignment/server/, Katoh et al. 2017)^37^ with ClustalW coloring. Both amino acids 229G and 2702D are highly conserved across diverse species, as well as their surrounding protein domains.

To investigate the role of genetic variation on regulatory importance in the context of super-enhancers, we aggregated rare non-coding variants across a set of 1,065 published MK-specific super-enhancers (***Supplementary Figure 5***)^33^. We found rare non-coding variants in a super-enhancer at the *PEAR1* locus was significantly associated with ADP-(P=2.4×10^−8^), epinephrine-(P-value=1.1×10^−7^) and collagen-(P-value=2.7×10^−5^) induced platelet aggregation. We observed, in marked contrast to our gene-based coding variant analyses, that the association signal in the *PEAR1* super-enhancer is driven by multiple rare variants in the region (***Supplementary Figure 6***).

In this first WGS study of platelet aggregation, we identify and replicate several novel loci contributing to trait variation. Among the 16 identified independent variants, 7 were associated with wide range of blood-related traits including CVD outcomes in a phenome-wide association study using UK Biobank data (***Supplementary Table 6***). While population genetic and genomic studies remain limited in size for platelet function traits, our results indicate the potential for novel discoveries made through sequencing approaches in deeply phenotyped individuals which may ultimately relate to CVD risk and treatment efficacy. This is underlined by data supporting the hypothesis that hyper-reactive platelets may predict future thromboses in both healthy, untreated individuals ^34^ and those who have already experienced thrombosis ^35,36^, as well as the fact that antiplatelet therapy remains standard of care in secondary prevention of atherothrombotic diseases such as myocardial infarction and stroke.

## Supporting information

Supplementary pictures

## Acknowledgements

Whole genome sequencing (WGS) for the Trans-Omics in Precision Medicine (TOPMed) program was supported by the National Heart, Lung and Blood Institute (NHLBI). WGS for GeneSTAR (Genetic Study of Atherosclerosis Risk) (phs001218) was performed at Macrogen, Illumina, and the Broad Institute of MIT and Harvard (HHSN268201500014C). WGS for the Old Order Amish (Genetics of Cardiometabolic Health in the Amish) (phs000956) was performed at the Broad Institute of MIT and Harvard (3R01HL121007-01S1). WGS for The Framingham Heart Study (Whole Genome Sequencing and Related Phenotypes in the Framingham Heart Study) (phs000974.v1.p1) was performed at the Broad Institute of MIT and Harvard (HHSN268201500014C). Centralized read mapping and genotype calling, along with variant quality metrics and filtering were provided by the TOPMed Informatics Research Center (3R01HL-117626-02S1). Phenotype harmonization, data management, sample-identity QC, and general study coordination, were provided by the TOPMed Data Coordinating Center (3R01HL-120393-02S1). We gratefully acknowledge the studies and participants who provided biological samples and data for TOPMed.

For the Old Order Amish this investigation was supported by National Institutes of Health grants U01 GM074518, U01 HL105198, R01 HL137922, R01 HL121007, and the University of Maryland Mid-Atlantic Nutrition and Obesity Research Center (P30 DK072488). GeneSTAR was supported by the National Institutes of Health/National Heart, Lung, and Blood Institute (U01 HL72518, HL087698, HL112064, HL11006, HL118356) and by a grant from the National Institutes of Health/National Center for Research Resources (M01-RR000052) to the Johns Hopkins General Clinical Research Center. The Framingham Heart Study is conducted and supported by the NHLBI in collaboration with Boston University (Contract No. N01-HC-25195), its contract with Affymetrix, Inc., for genome-wide genotyping services (Contract No. N02-HL-6–4278 and Contract No. HHSN268201500001I). MHC, BATC and ADJ were supported by NHLBI Intramural funding. The Caerphilly Prospective study was undertaken by the former MRC Epidemiology Unit (South Wales) and was funded by the Medical Research Council of the UK. The data archive is maintained by the School of Social and Community Medicine, University of Bristol.This study makes use of data generated by the BLUEPRINT Consortium. A full list of the investigators who contributed to the generation of the data is available from www.blueprint-epigenome.eu. Funding for the project was provided by the European Union’s Seventh Framework Programme (FP7/2007-2013) under grant agreement no 282510 BLUEPRINT. The views expressed in this manuscript are those of the authors and do not necessarily represent the views of the National Heart, Lung, and Blood Institute; the National Institutes of Health; or the U.S. Department of Health and Human Services.

## Author contribution

A.R.K., B.A.T.R, M.-H.C, J.P.L, R.A.M, A.D.J led the study. A.R.K wrote the first draft of manuscript with contribution and editing from B.A.T.R, M.-H.C, L.R.Y, M.A.T, J.A.B, L.C.B, N.F, J.P.L, R.A.M and A.D.J. Genome wide analyses were performed by A.R.K., B.A.T.R, B.J.G, L.R.Y, M.-H.C and K.R. RNA-sequencing and eQTL analyses were performed by R.A.M, K.K, M.A.T, L.R.Y, A.R.K., K.K. and K.I. Imputation of genomic data and replication analyses were performed by M.-H.C, B.A.T.R, L.R.Y, B.J.G, A.P, L.A.C, and M.H.K. L.R.Y, N.F, L.C.B and R.A.M were involved in the guidance, collection and analysis for Genetic Study of Atherosclerosis Risk (GeneSTAR) phenotype data. B.J. G., K.R., B.D.M, J.P.L, and A.R.S were involved in the guidance, collection and analysis for Older Order Amish Study (OAA) phenotype data. B.A.T.R., M.-H.C and A.D.J. were involved in the guidance and analysis for Framingham Heart Study (FHS) phenotype data. A.D.J. funded genotyping of the CaPs cohort. M.-H.C. performed genotype QC, calling and imputation of the CaPs cohort with input from A.D.J. B.A.T.R., M.-H.C. and A.D.J. conducted CaPs genotype-phenotype analyses.

