## Supplementary pictures for "Genome Sequencing Unveils a New Regulatory Landscape of Platelet Reactivity"

Table of contents:

Supplementary Figure1 ------------------------------------------------------------------- Page 3
Supplementary Figure2 ------------------------------------------------------------------- Page 10
Supplementary Figure3 ------------------------------------------------------------------- Page 13
Supplementary Figure4 ------------------------------------------------------------------- Page 14
Supplementary Figure5 ------------------------------------------------------------------- Page 15
Supplementary Figure6 ------------------------------------------------------------------- Page 16

**Supplementary Figure 1.** Manhattan plots and Quantile-Quantile (QQ) plots of platelet aggregation in response to different doses of Epinephrine, ADP and Collagen as described in Supplementary Table 1. Genome-wide association study for platelet aggregation in 3,855 individuals. P-values, expressed as -log 10(P), are plotted according to physical genomic locations by chromosome. Loci passing genome wide significance ($5 \times 10^{-8}$) are marked by red dots. Locus names represent the nearest annotated gene. The blue horizontal line indicates a P-value threshold of $1 \times 10^{-6}$ corresponding to suggestive significance threshold. The red horizontal line indicates P-value threshold of $5 \times 10^{-8}$, corresponding to genome-wide significance.

ADP low 1

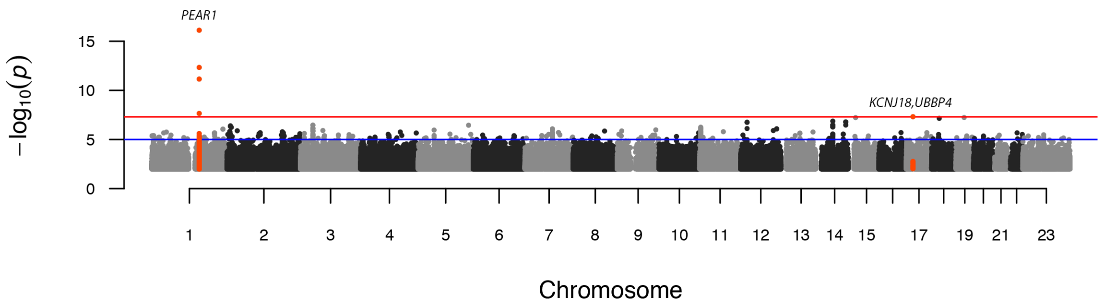


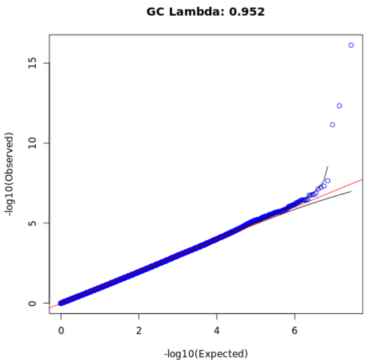


ADP low 2

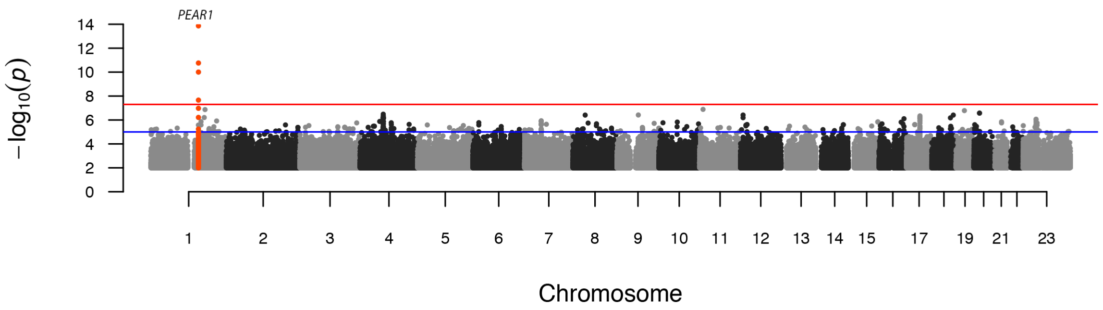


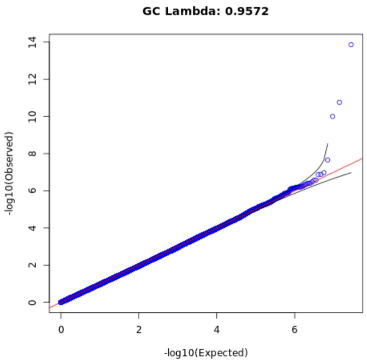

#### ADP low 3

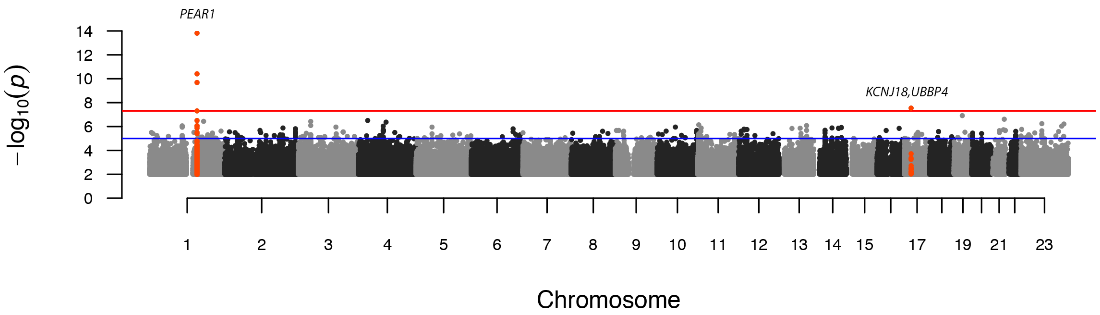


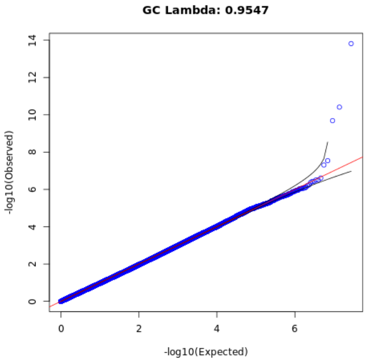


#### ADP high 1

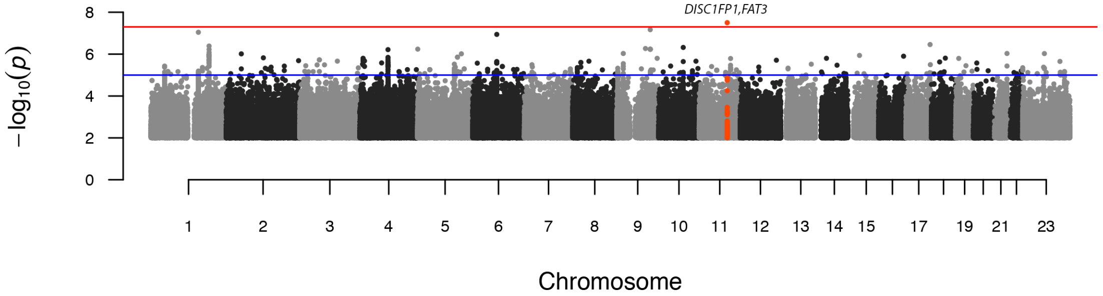


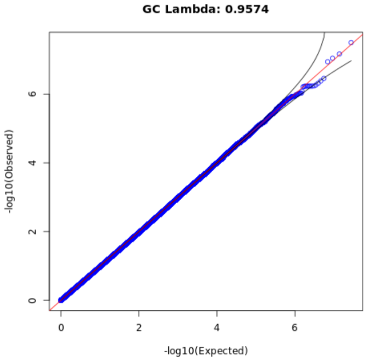


#### ADP high 2

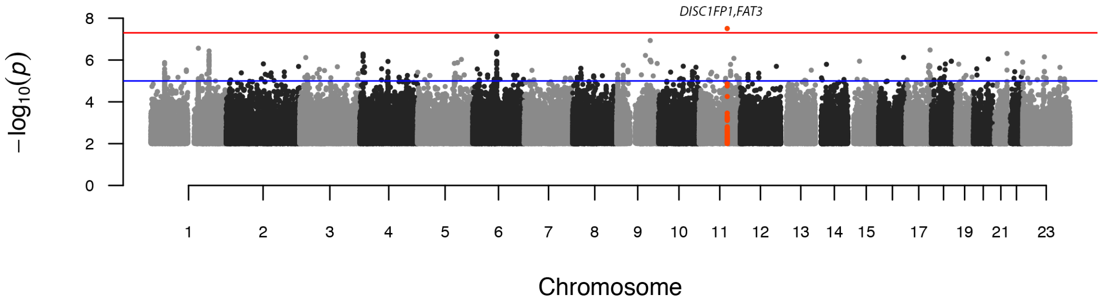


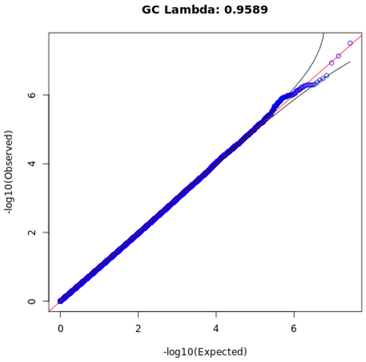

ADP high 3

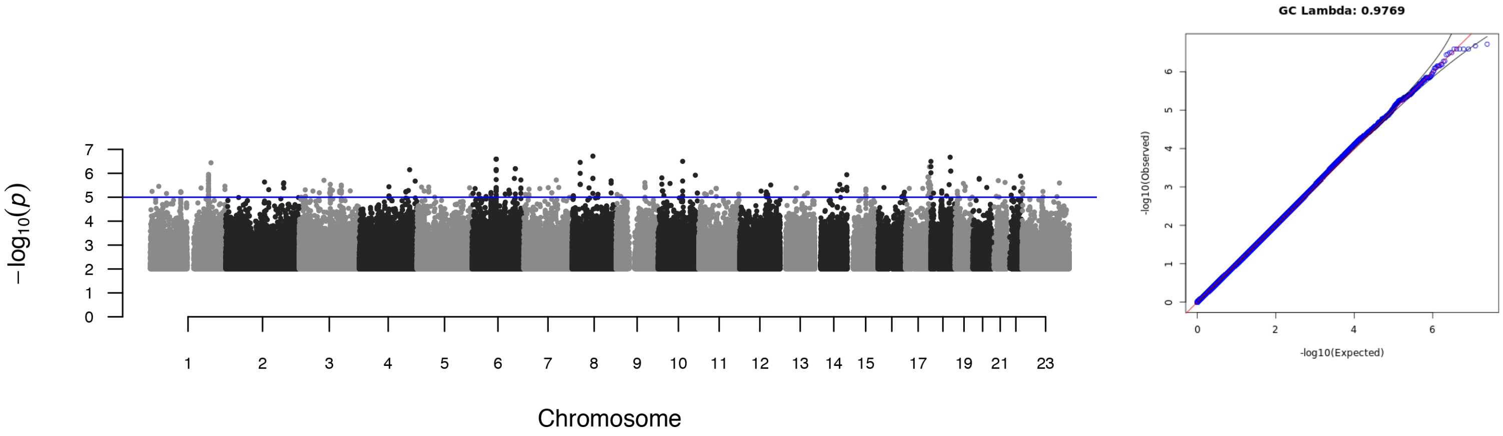


ADP high 4

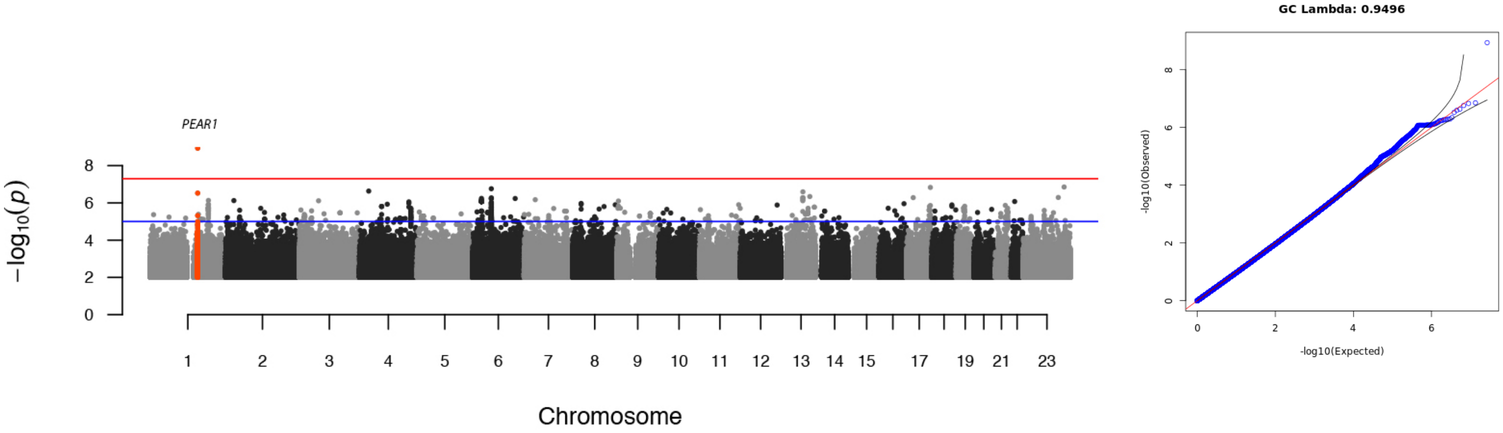


Collagen low 1

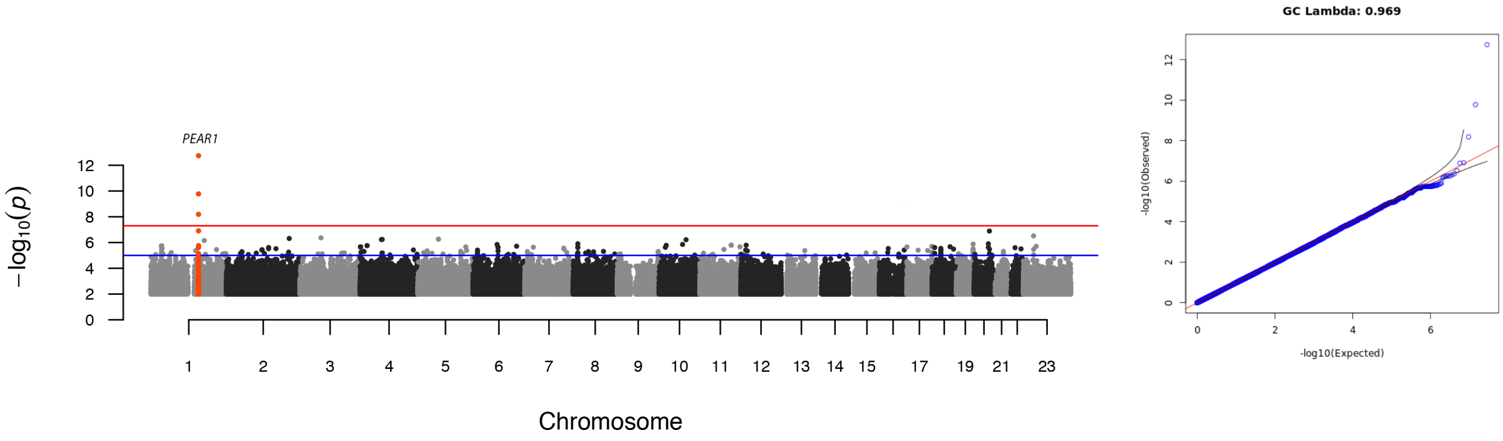

Collagen low 2

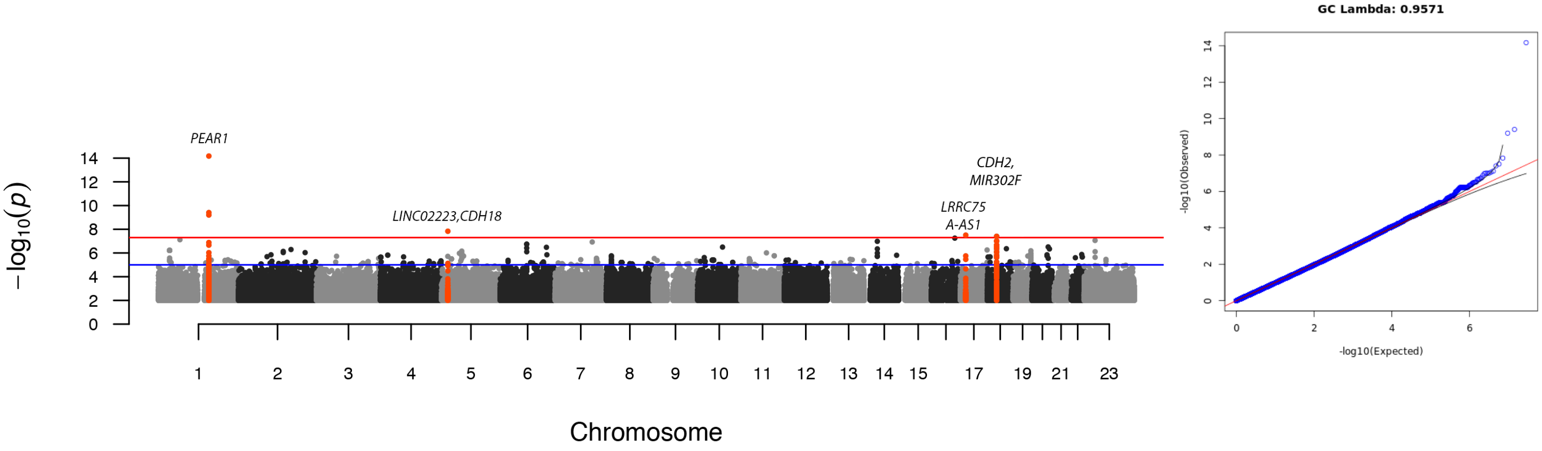


Collagen high 1

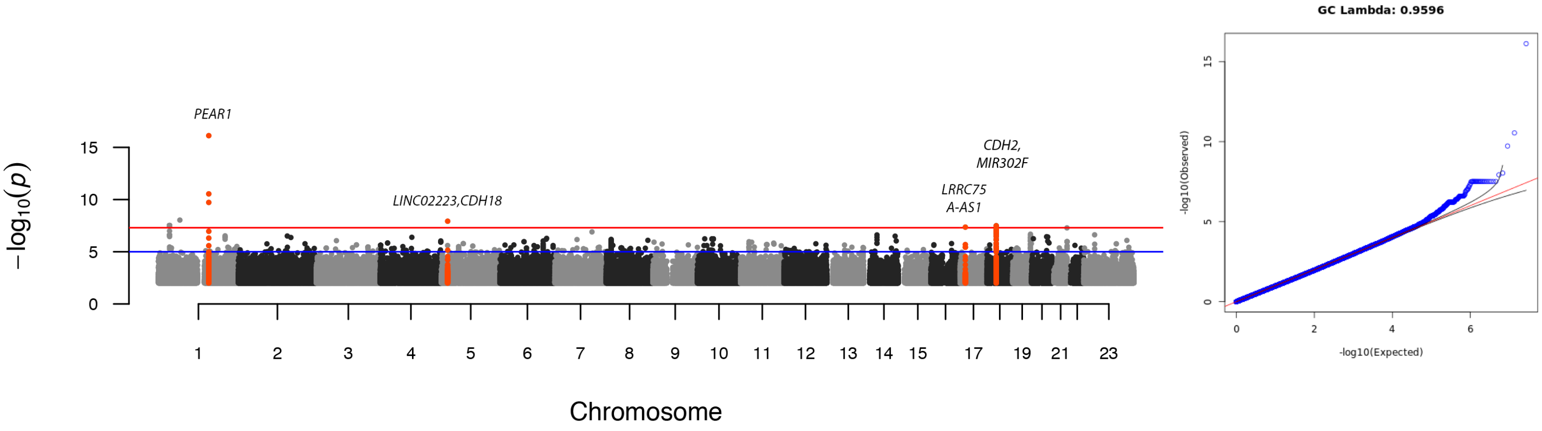


Collagen high 2

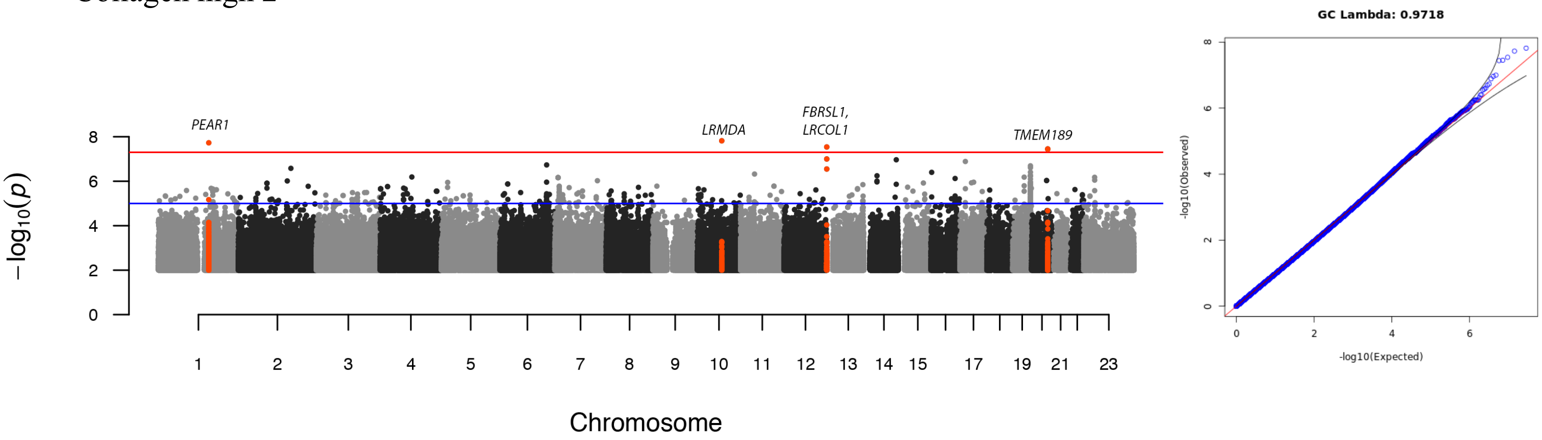

Epinephrine low 1

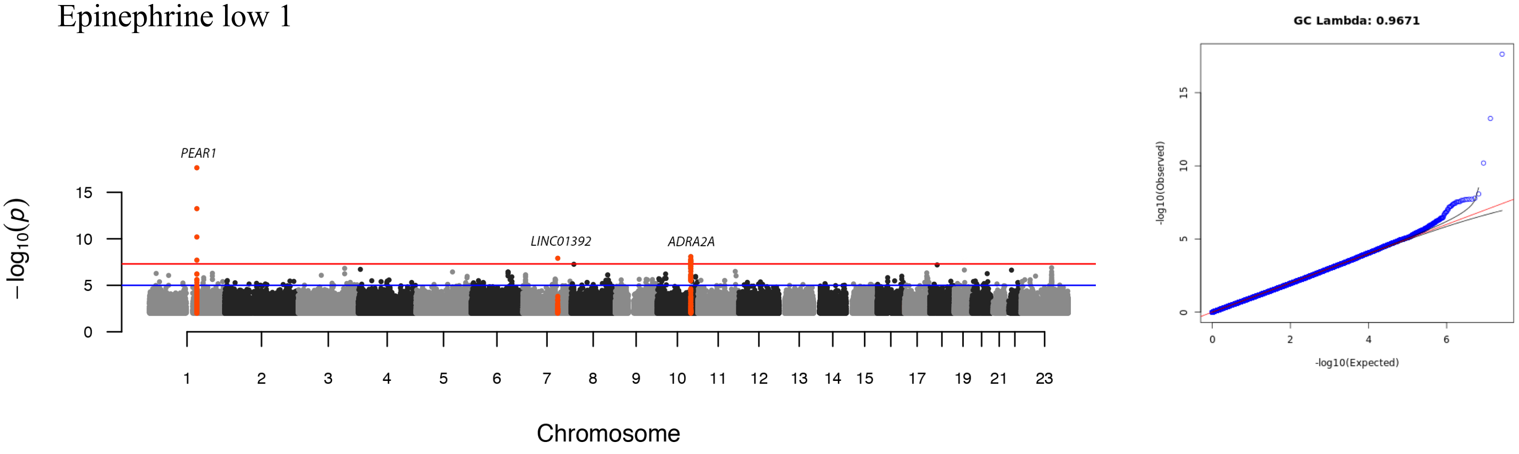


Epinephrine low 2

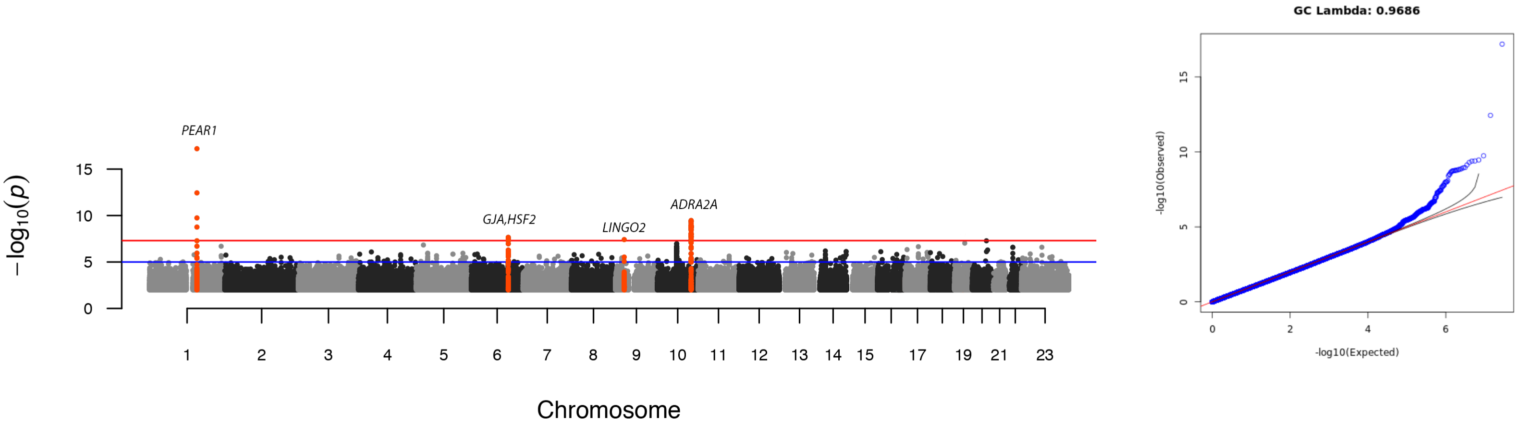


Epinephrine low 3

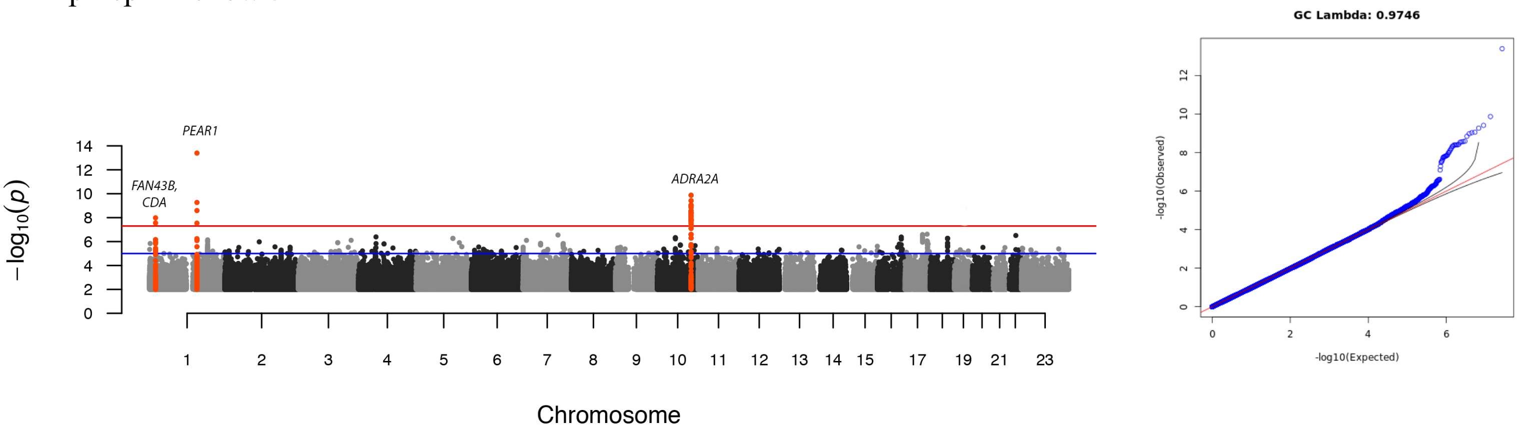

#### Epinephrine low 4

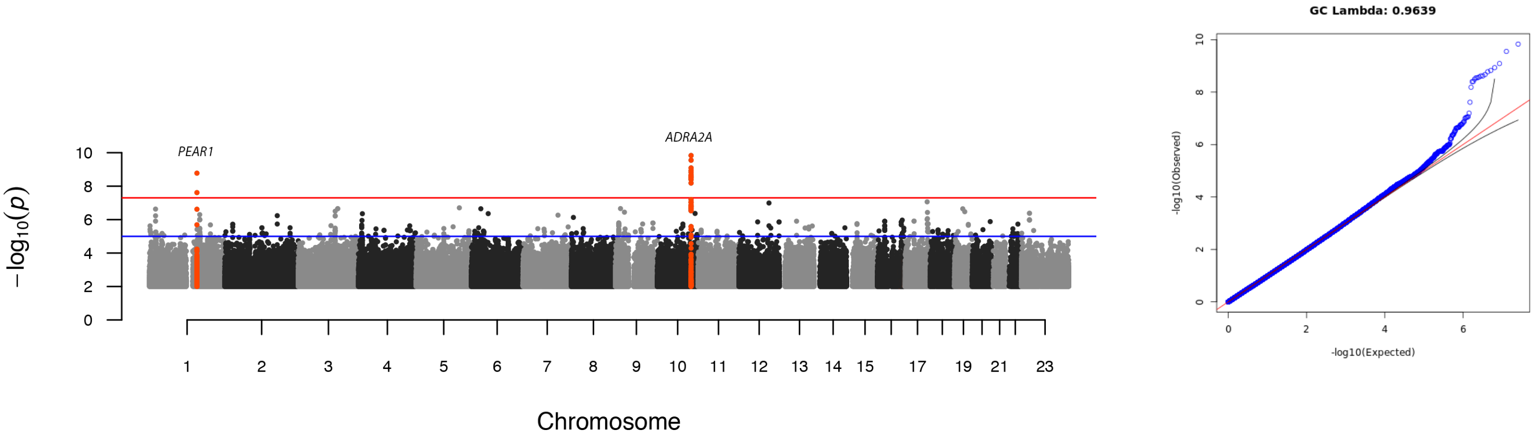


#### Epinephrine low 5

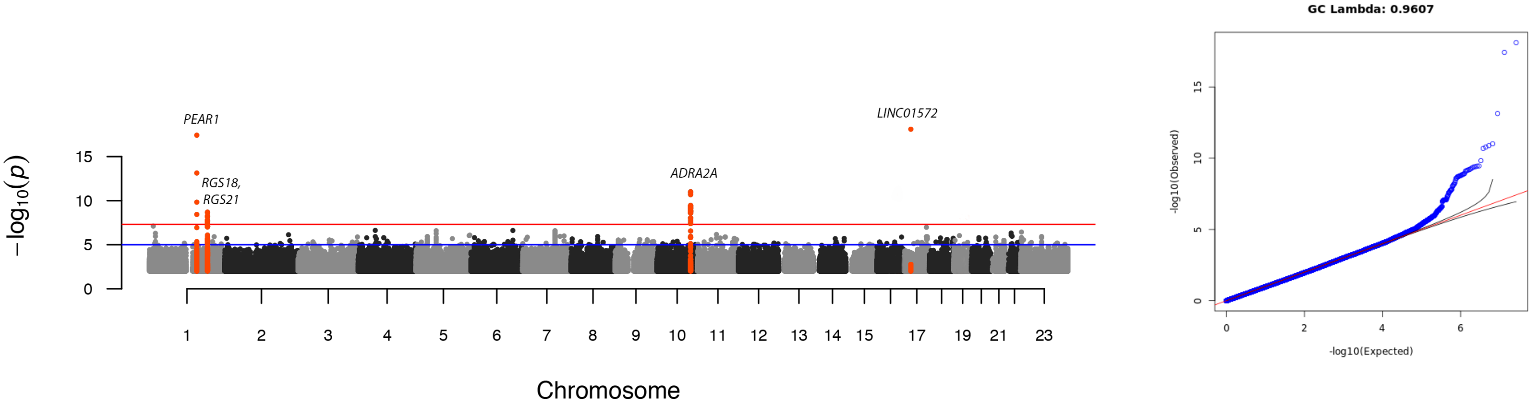


#### Epinephrine high 1

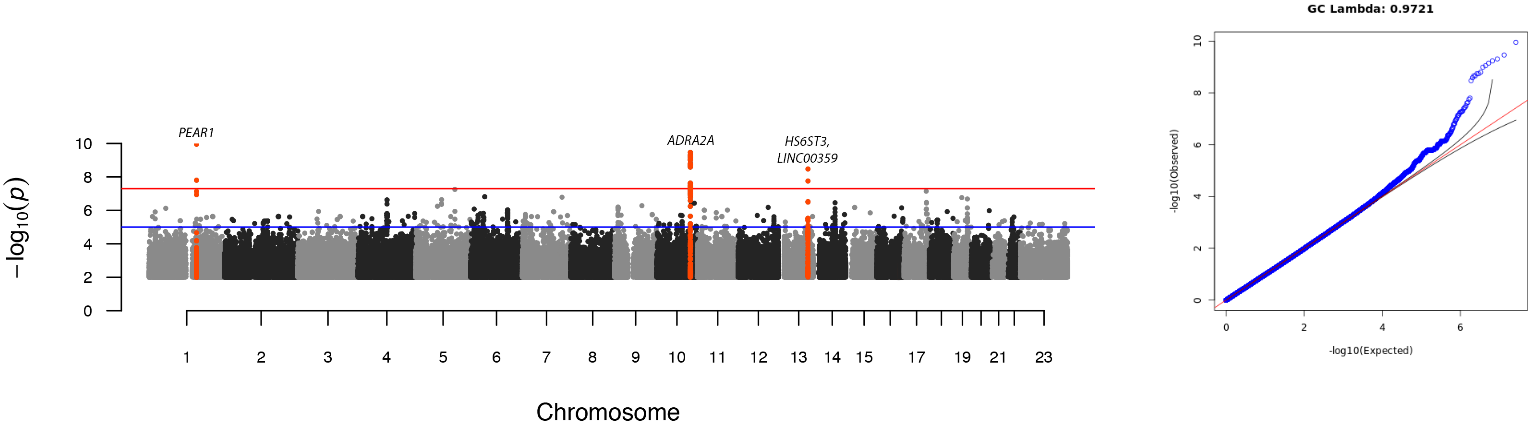

### Epinephrine high 2

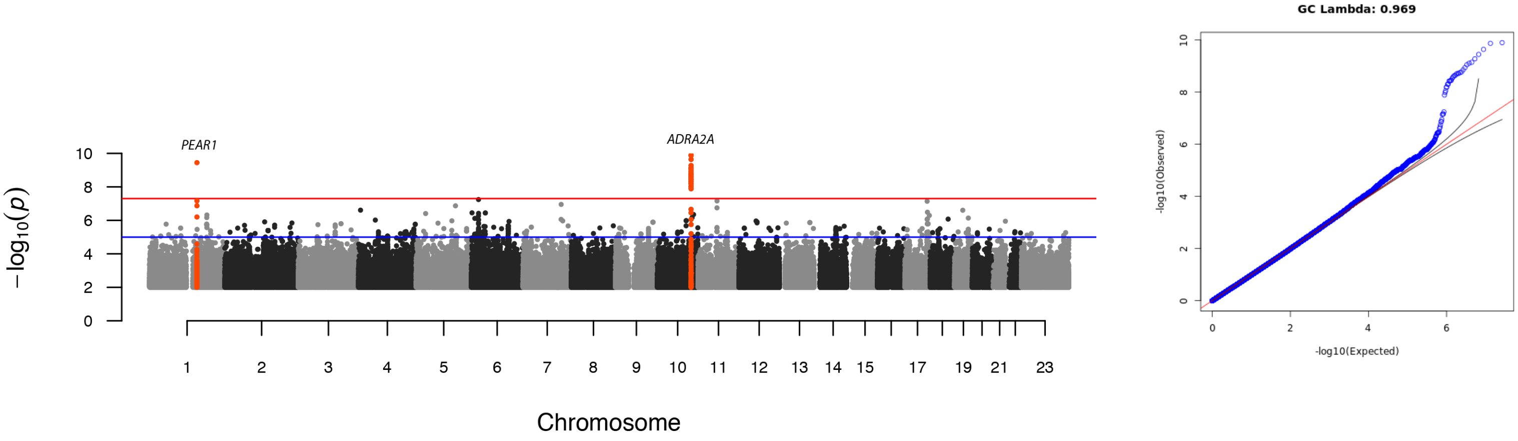


### Epinephrine high 3

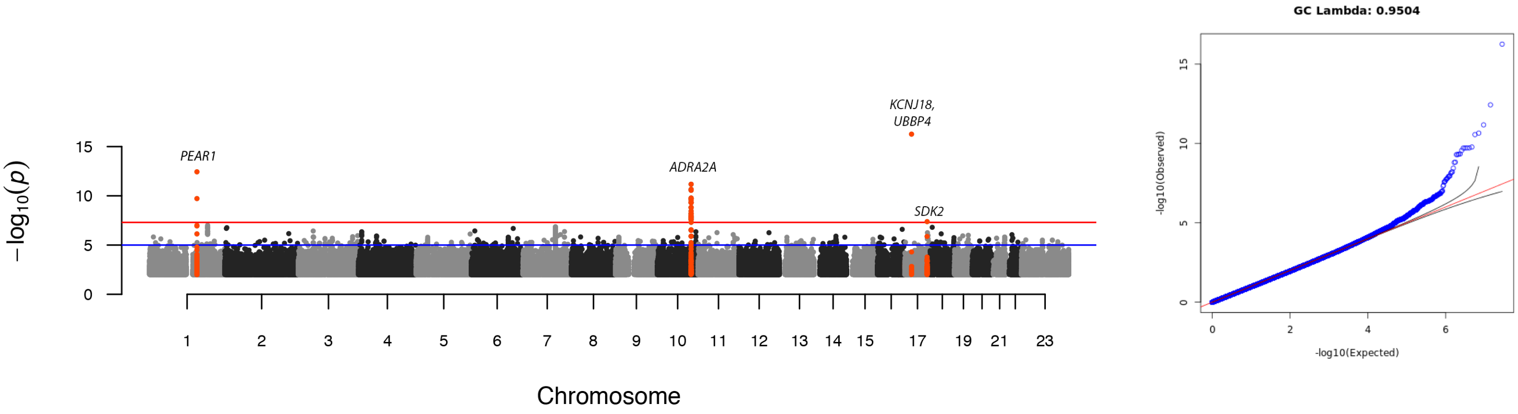

**Supplementary Figure 2.** Locus zoom plots of independent loci associated with platelet aggregation in response to ADP(2a), Collagen(2b) and Epinephrine(2c).

2a-ADP

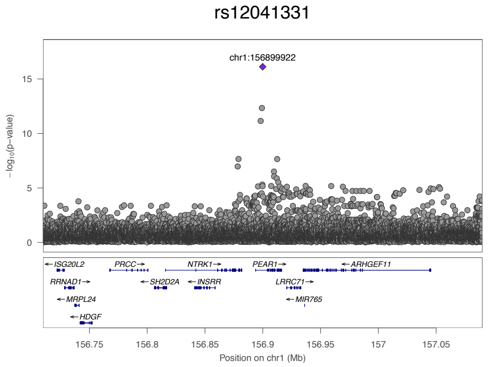


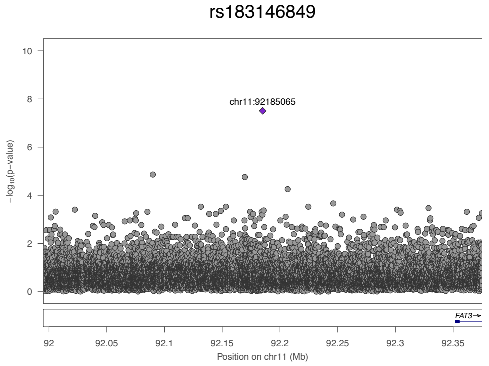


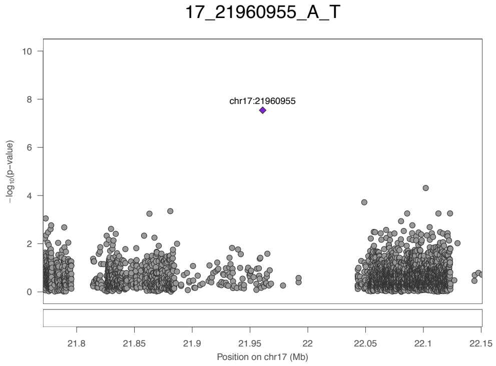

### 2b-Collagen

rs12041331

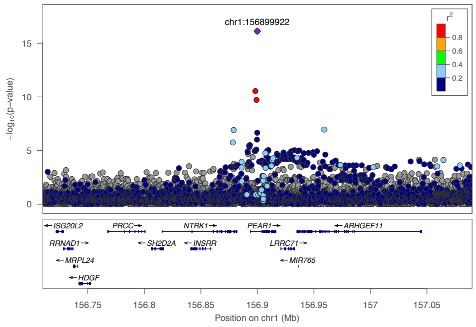


rs140148392

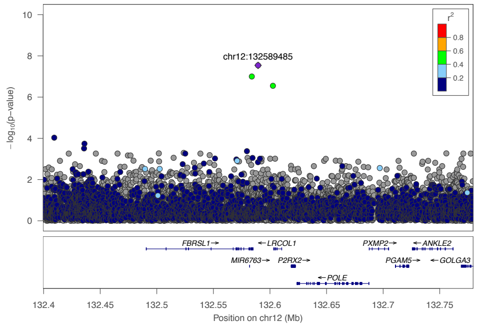


rs142001088

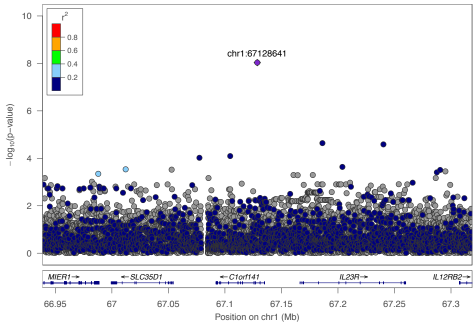


rs575524466

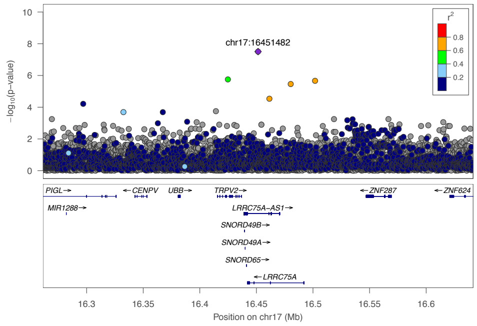


rs112157462

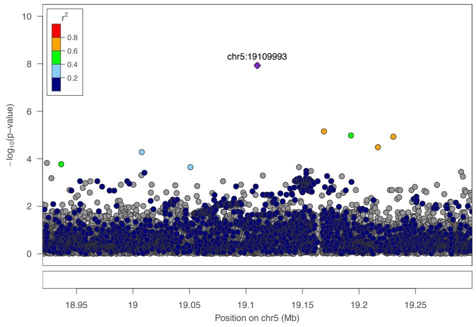


rs138845468

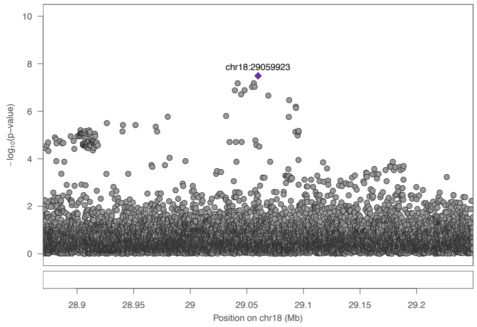


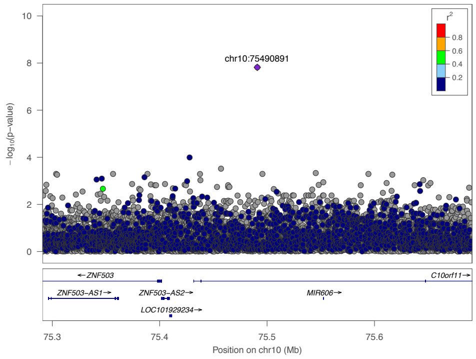


rs542707094

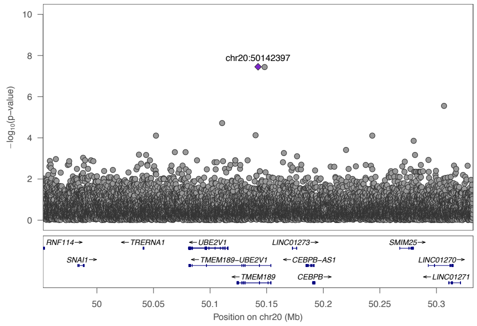

### 2c-Epinephrine

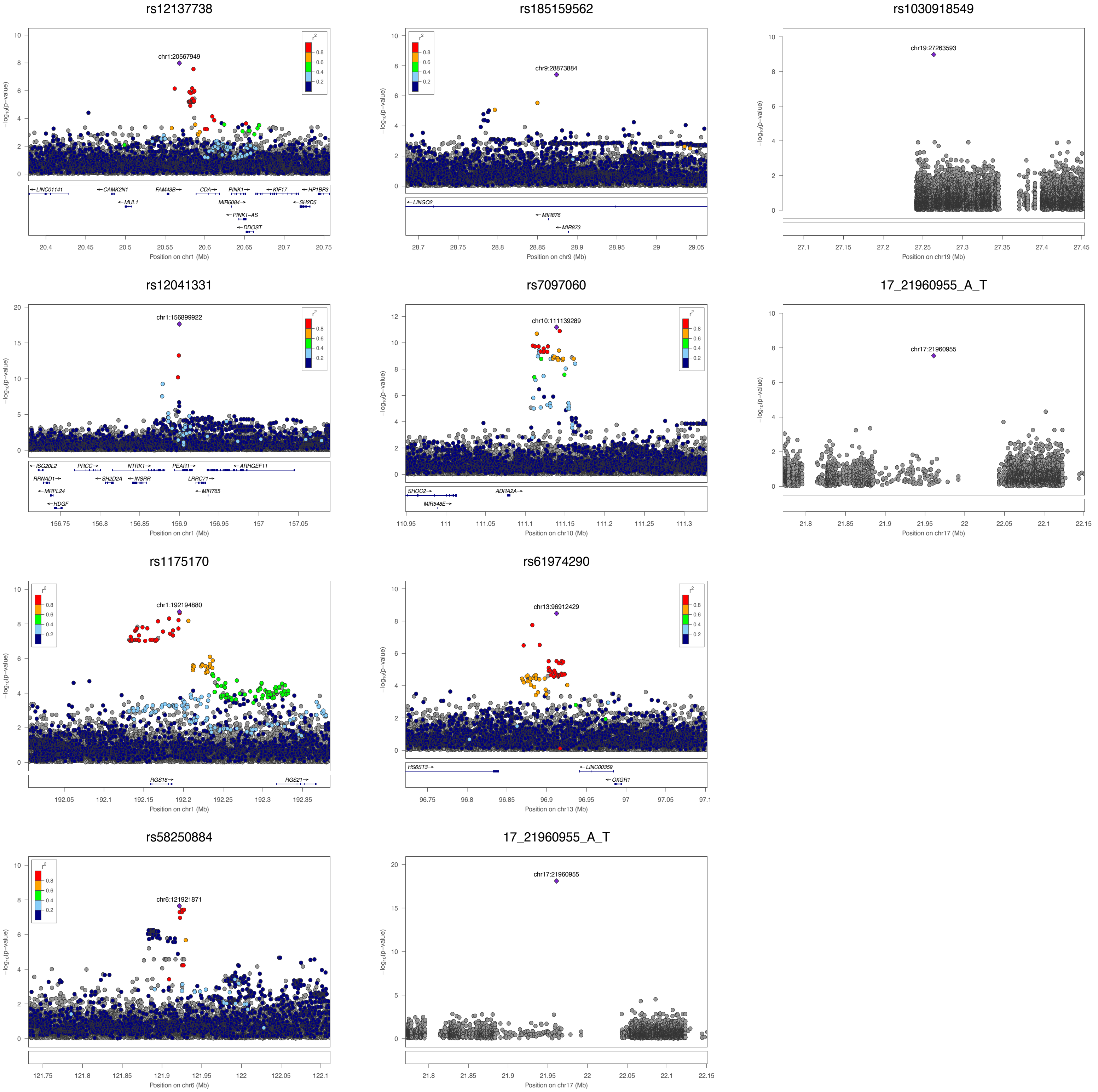

**Supplementary Figure 3:** QQ plots of SKAT analyses of rare deleterious coding variants with ADP, Collagen and Epinephrine induced platelet aggregation.

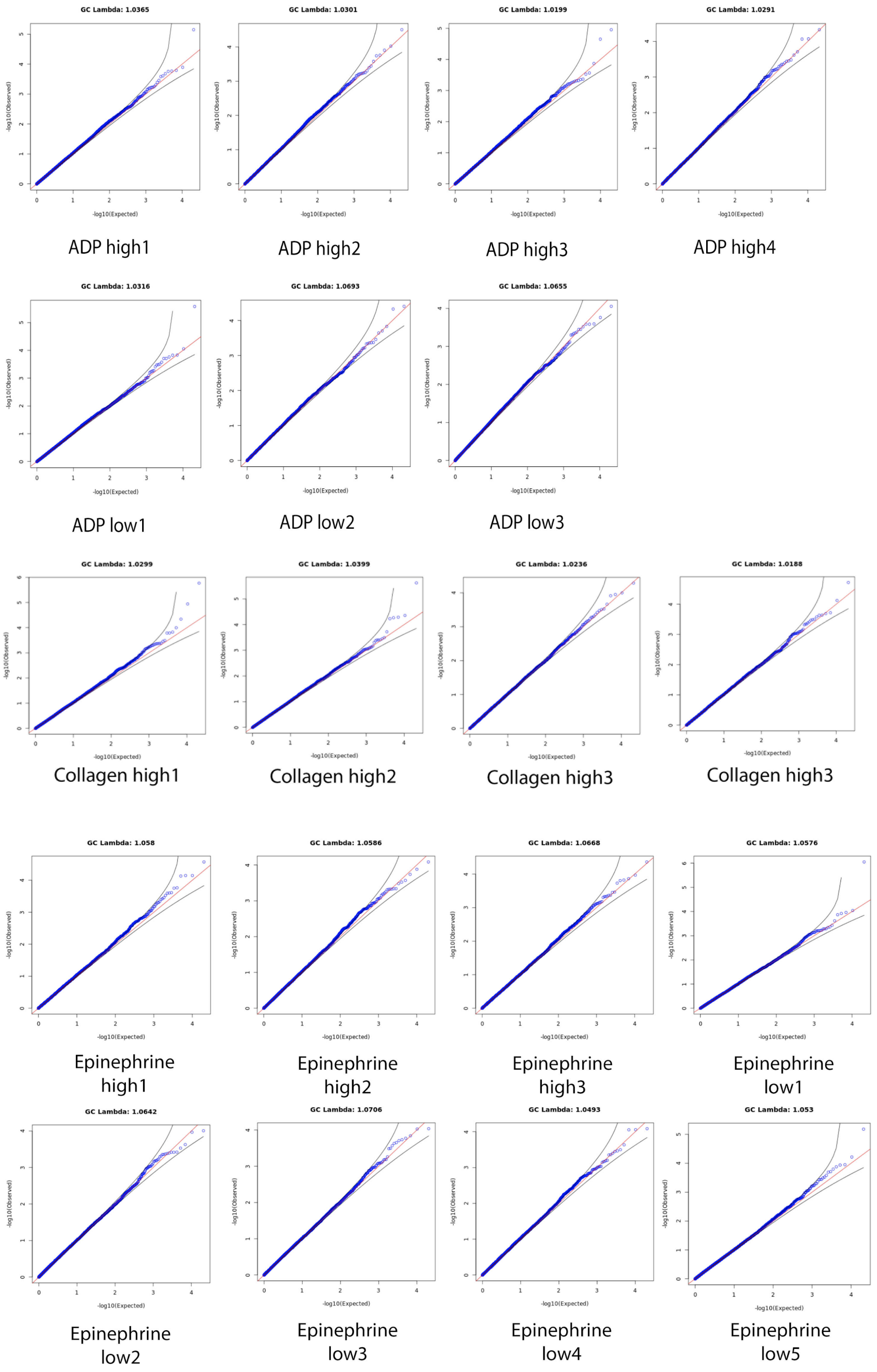

**Supplementary Figure 4:** Leave-one-out technique to identify the variants contributing the most to SKAT p-value of SVEP1(4a), BCO1(4b), NELFA(4c) and IDH3A(4d) were associated with platelet aggregation after Bonferroni correction (0.05 / 17744 = 2.819E$^{-6}$). x-axis represents the variants in the gene set, orange dots represent the –Log10(p-value) of gene set when the variant was left out and blue bars represent the minor allele frequency of the variants.

**4a**

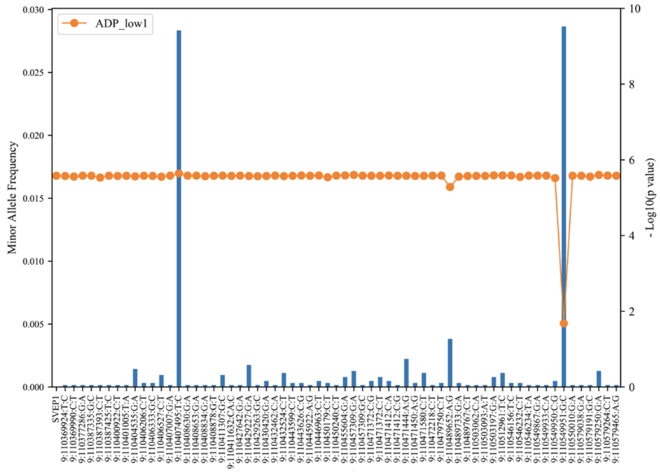


**4b**

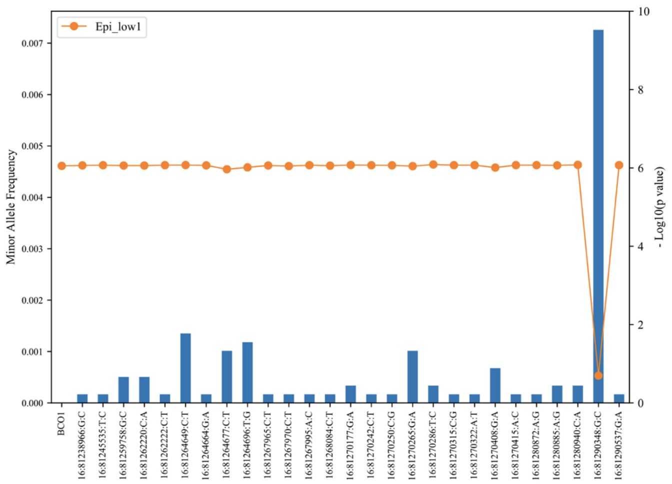


**4c**

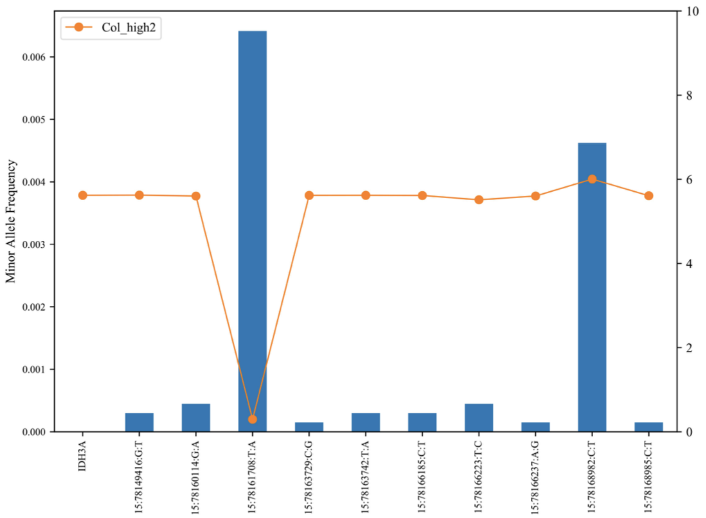


**4d**

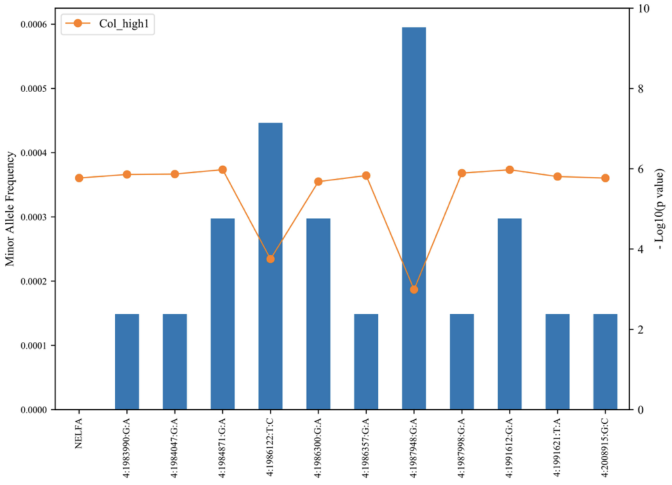

**Supplementary Figure 5:** QQ plots of SKAT analyses of rare non-coding variants in megakaryocyte specific super-enhancers with ADP, Collagen and Epinephrine induced platelet aggregation.

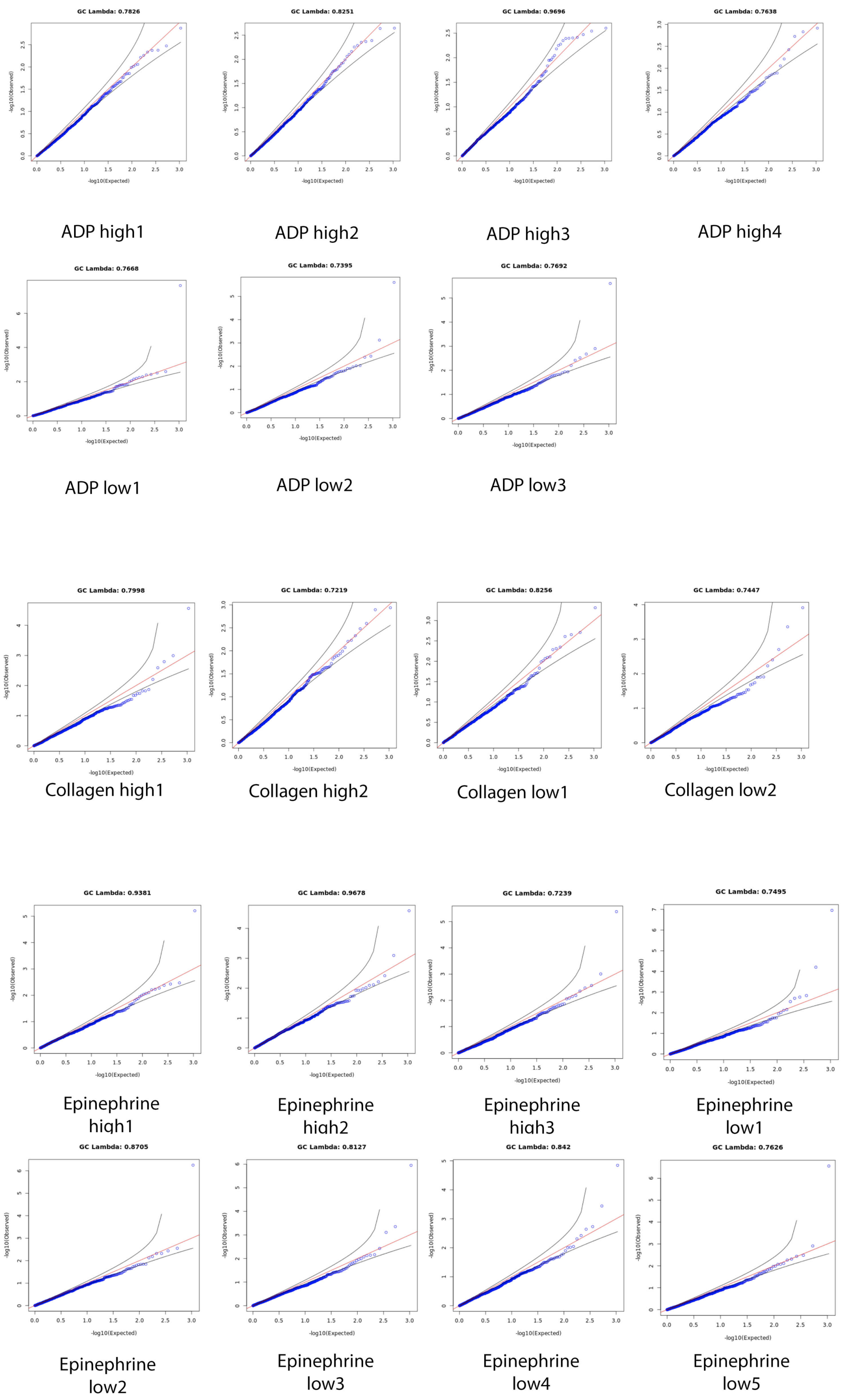

**Supplementary Figure 6:** Leave-one-out technique to identify the variants contributing the most to SKAT p-value of the super enhancer located at *PEAR1* locus. Blue dots represent the -Log10(p-value) of aggregated non-coding variants in *PEAR1* locus when the variant was left out.

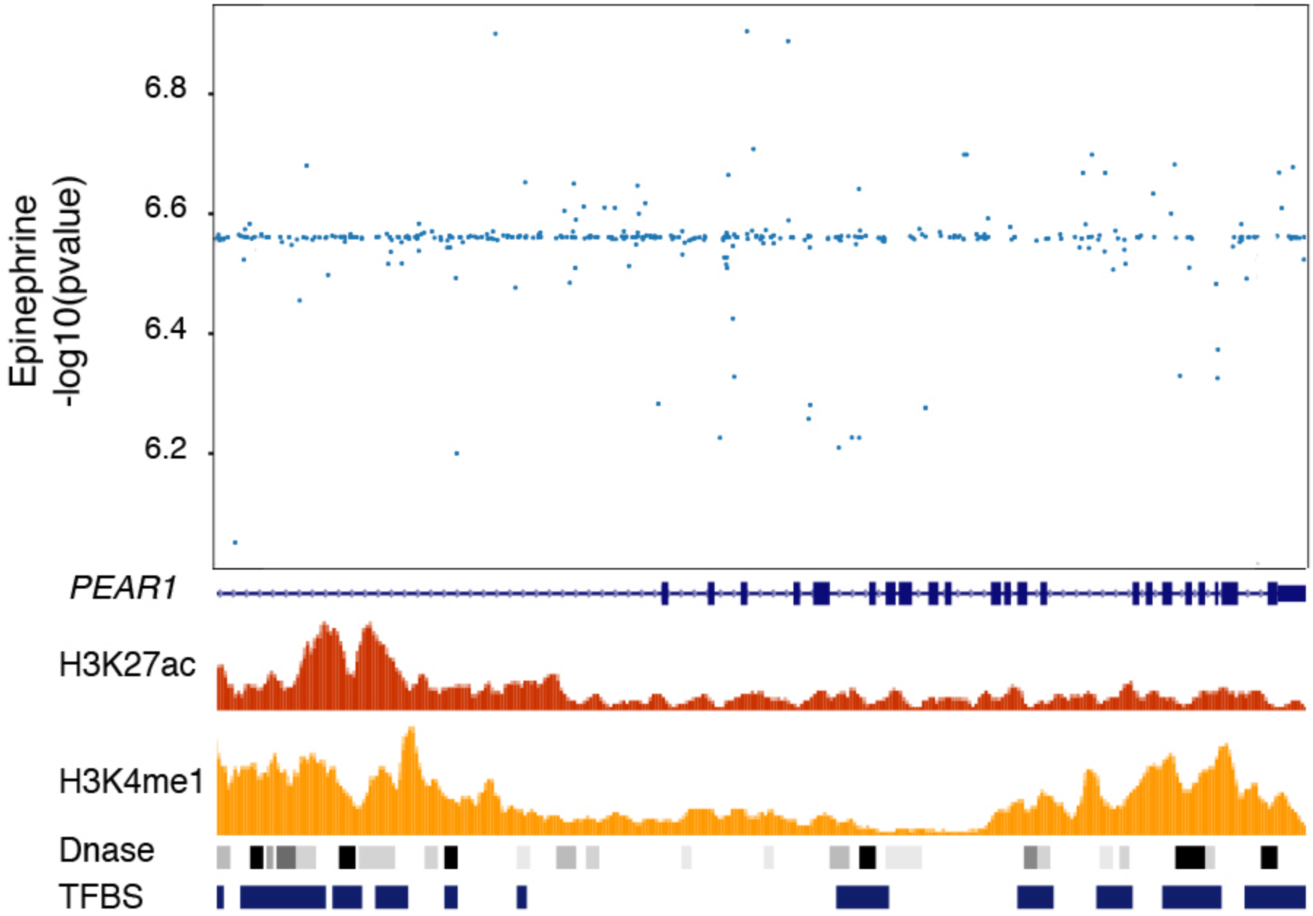
