## Supplementary tables for "Genome Sequencing Unveils a New Regulatory Landscape of Platelet Reactivity"

**^++^** *https://www.nhlbiwgs.org/topmed-banner-authorship*

1 Division of Cardiology, Johns Hopkins University School of Medicine, Baltimore

2 GeneSTAR Research Program, Johns Hopkins University School of Medicine, Baltimore

3 Division of Intramural Research, Population Sciences Branch, National Heart, Lung and Blood Institute, Bethesda

4 The Framingham Heart Study, Framingham

5 Division of General Internal Medicine, Johns Hopkins University School of Medicine, Baltimore

6 Division of Endocrinology, Diabetes, and Nutrition, University of Maryland School of Medicine, Baltimore

7 Program in Personalized and Genomic Medicine, University of Maryland School of Medicine, Baltimore, Baltimore

8 Cardiovascular Health Research Unit, University of Washington School of Medicine, Seattle

9 Biostatistics and Bioinformatics, Oncology, Sidney Kimmel Comprehensive Cancer Center, Johns Hopkins University School of Medicine, Baltimore

10 Division of Allergy and Clinical Immunology, Johns Hopkins University School of Medicine, Baltimore

11 Department of Biostatistics, University of North Carolina, Chapel Hill

12 Department of Biostatistics, School of Public Health, Boston University, Boston

13 Department of Anesthesiology and Critical Care Medicine, Johns Hopkins University School of Medicine, Baltimore

14 Bloomberg School of Public Health, Biostatistics, Johns Hopkins University, Baltimore

Address Correspondence to: **Joshua P. Lewis, PhD,**, 410-706-5087; **Rasika A Mathias, ScD**,, 410-550-2487; **Andrew D. Johnson, PhD**,, 508-663-4082.

Table of contents:

Supplementary Table 1 ------------------------------------------------------------------------- Page 2

Supplementary Table 2 ------------------------------------------------------------------------- Page 3

Supplementary Table 3a ------------------------------------------------------------------------ Page 4-5

Supplementary Table 3b ------------------------------------------------------------------------ Page 6

Supplementary Table 4 -------------------------------------------------------------------------- Page 7

Supplementary Table 5 -------------------------------------------------------------------------- Page 8

Supplementary Table 6 -------------------------------------------------------------------------- Page 9

Supplementary Table 7 -------------------------------------------------------------------------- Page 10

Supplementary Table 8 -------------------------------------------------------------------------- Page 11

Supplementary Table 9 -------------------------------------------------------------------------- Page 12

Supplementary Table 10 ------------------------------------------------------------------------ Page 13

**Supplementary Table 1**. Platelet aggregation results were harmonized across the cohorts. * indicates Maximum aggregation to ADP/Epinephrine and ** indicates Lag time to Collagen.

| **Phenotype** | **Agonist** | **Framingham** | **GeneSTAR** | **Amish** | **Sample size** |
| --- | --- | --- | --- | --- | --- |
| adp_low1 | ADP | * 1μM | * 2μM | * 2μM | 3140 |
| adp_low2 | ADP | * 3μM | * 2μM | * 2μM | 3229 |
| adp_low3 | ADP | Threshold dose for >50% aggregation | * 2μM | * 2μM | 3014 |
| adp_high1 | ADP | * 5μM | * 10μM | * 5μM | 2799 |
| adp_high2 | ADP | * 5μM | * 10μM | * 10μM | 2799 |
| adp_high3 | ADP | * 10μM | * 10μM | * 10μM | 1967 |
| adp_high4 | ADP | Threshold dose for >50% aggregation | * 10μM | * 10μM | 3147 |
| epi_low1 | Epinephrine | * 0.5μM | * 2μM | * 10μM | 2962 |
| epi_low2 | Epinephrine | * 1μM | * 2μM | * 10μM | 3027 |
| epi_low3 | Epinephrine | * 3μM | * 2μM | * 10μM | 2486 |
| epi_low4 | Epinephrine | * 3μM | * 10μM | * 10μM | 2488 |
| epi_low5 | Epinephrine | Threshold dose for >50% aggregation | * 2μM | * 10μM | 3152 |
| epi_high1 | Epinephrine | * 5μM | * 10μM | * 10μM | 2098 |
| epi_high2 | Epinephrine | * 10μM | * 10μM | * 10μM | 2141 |
| epi_high3 | Epinephrine | Threshold dose for >50% aggregation | * 10μM | * 10μM | 3154 |
| col_low1 | Collagen | ** 190 μg/mL | ** 1μg/mL | ** 1μg/mL | 3354 |
| col_low2 | Collagen | ** 190 μg/mL | ** 2μg/mL | ** 2μg/mL | 3364 |
| col_high1 | Collagen | ** 190 μg/mL | ** 5μg/mL | ** 5μg/mL | 3361 |
| col_high2 | Collagen | ** 190 μg/mL | ** 10μg/mL | ** 10μg/mL | 3352 |

**Supplementary Table 2.** Demographics for participants included in genome-wide association analyses. Values given are % or mean ± 1 SD. FHS=Framingham Heart Study, Amish=Old Order Amish Study, GS=GeneSTAR, EA=European American, AA=African American.

| **Characteristic** | **FHS**  **n=1,981** | **Amish**  **n=235** | **GS EA**  **n=909** | **GS AA**  **n=730** |
| --- | --- | --- | --- | --- |
| Male, % | 47.15 | 48.94 | 44.11 | 36.71 |
| Diabetes, % | 6.30 | 0.43 | 5.19 | 11.96 |
| Hypertension, % | 35.34 | 5.96 | 24.67 | 39.25 |
| Smoking, % | 17.47 | 8.94 | 20.59 | 31.85 |
| Cardiovascular disease, % | 6.11 | 2.55 | 1.10 | 1.78 |
| Aspirin response, % | 15.49 | 0 | 0 | 0 |
| Age, years | 55.76 ± 9.2 | 46.73 ± 13.6 | 44.5 ± 13.2 | 43.4 ± 12.4 |
| Body mass index, kg/m2 | 27.52 ± 4.9 | 26.9 ± 4.4 | 28.7 ± 6.4 | 31.8 ± 8.1 |
| LDL cholesterol, mg/dl | 127.4 ± 32.8 | 144.6 ± 44.8 | 124.8 ± 37.4 | 120.6 ± 38.9 |
| Fibrinogen, mg/dl | 308.0 ± 56.2 | 281.9 ± 57.8 | 374.3 ± 113 | 417.3 ± 122 |
| Maximal aggregation to low ADP doses (%) | 19.0 ± 21.5 (1uM)  67.8 ± 24.7(3uM) | 41.5 ± 22.5 (2uM) | 44.6 ± 26.2 (2uM) | 41.8 ± 28.6 (2uM) |
| Maximal aggregation to high ADP doses (%) | 77.4 ± 18.9 (5uM)  78.0 ± 19.5 (10uM) | 61.5 ± 18.4 (5uM)  67.5 ± 13.5 (10uM) | 79.4 ± 13.3 (10uM) | 77.0 ± 17.3 (10uM) |
| Threshold concentration (EC50) for 50% response to ADP (*u*M) | 3.27 ± 1.5 | NA | NA | NA |
| Maximal aggregation to low epinephrine doses (% ) | 51.2 ± 31.0 (0.5uM)  58.2 ± 31.1 (1uM)  66.0 ± 28.4 (3uM) | 60.2 ± 27.1 (10uM) | 56.1 ± 33.2 (2uM) | 51.6 ± 36.0 (2uM) |
| Maximal aggregation to high epinephrine doses (%) | 48.1 ± 28.6 (5uM)  34.8 ± 24.5 (10uM) | 60.2 ± 27.1 (10uM) | 71.8 ± 27.2 (10uM) | 63.5 ± 34.1 (10uM) |
| Threshold concentration (EC50) for 50% response to epinephrine (*u*M) | 1.92 ± 2.9 | NA | NA | NA |
| Lag time to low collagen doses (seconds) | 87.0 ± 25.2 (190ug/ml) | 56.8 ± 24.9 (1ug/ml)  47.9 ± 17.5 (2ug/ml) | 147.8 ± 87.6 (1ug/ml)  99.9 ± 57.5 (2ug/ml) | 158.1 ± 90.4 (1ug/ml)  111.5 ± 67.3 (2ug/ml) |
| Lag time to high collagen doses (seconds) | 87.0 ± 25.2 (190ug/ml) | 37.9 ± 12.1 (5ug/ml)  32.6 ± 10.5 (10ug/ml) | 68.8 ± 32.1 (5ug/ml)  60.5 ± 23.3 (10ug/ml) | 77.4 ± 40.8 (5ug/ml)  64.1 ± 26.4 (10ug/ml) |

**Supplementary Table 3a-** Discovery and Replication results by 16 Discovered Loci and phenotypes. Replication was performed as a sample-size weighted meta-analysis of up to N=2,009 individuals in order of FHS, GeneSTAR European Americans, GeneSTAR African Americans and OOA for direction of effect. An imputation Rsq filter of 0.7 was applied in the meta-analysis. Replication p-value was computed using one-sided test.

|  |  |  | **DISCOVERY** | | | | | **REPLICATION** | | | |
| --- | --- | --- | --- | --- | --- | --- | --- | --- | --- | --- | --- |
| **snpID hg38** | **ref/alt** | **Phenotype** | **N** | **MAF** | **beta** | **se** | **pval** | **Direction** | **N** | **Z-score** | **pval** |
| 1:20567949 | A/T | epi_low3 | 2486 | 0.076 | 0.306 | 0.053 | 1.04E-08 | -+++ | 1323 | 0.834 | 0.202 |
| 1:67128641 | C/T | col_high1 | 3361 | 0.020 | -0.503 | 0.088 | 9.25E-09 | +-?? | 1514 | 1.118 | 0.868 |
| 1:156899922 | G/A | epi_low1 | 2962 | 0.140 | -0.358 | 0.041 | 2.31E-18 | ---- | 1689 | -5.610 | 1.01E-08 |
|  |  | epi_low5 | 3152 | 0.139 | -0.345 | 0.040 | 3.72E-18 | ---- | 1855 | -6.179 | 3.23E-10 |
|  |  | epi_low2 | 3027 | 0.142 | -0.344 | 0.040 | 6.41E-18 | ---- | 1764 | -6.029 | 8.25E-10 |
|  |  | col_high1 | 3361 | 0.137 | 0.317 | 0.038 | 7.58E-17 | ++++ | 2008 | 5.076 | 1.93E-07 |
|  |  | adp_low1 | 3140 | 0.138 | -0.329 | 0.039 | 7.61E-17 | ---- | 1925 | -5.609 | 1.02E-08 |
|  |  | col_low2 | 3364 | 0.137 | 0.296 | 0.038 | 6.69E-15 | ++++ | 2009 | 5.146 | 1.33E-07 |
|  |  | adp_low2 | 3229 | 0.138 | -0.298 | 0.039 | 1.36E-14 | ---- | 2007 | -5.270 | 6.82E-08 |
|  |  | adp_low3 | 3014 | 0.140 | -0.310 | 0.040 | 1.52E-14 | ---- | 1837 | -5.748 | 4.52E-09 |
|  |  | epi_low3 | 2486 | 0.162 | -0.315 | 0.042 | 4.01E-14 | ---- | 1323 | -4.277 | 9.47E-06 |
|  |  | col_low1 | 3354 | 0.137 | 0.279 | 0.038 | 1.81E-13 | ++++ | 2009 | 5.883 | 2.01E-09 |
|  |  | epi_high3 | 3154 | 0.139 | -0.289 | 0.040 | 3.65E-13 | ---- | 1856 | -5.982 | 1.10E-09 |
|  |  | epi_high1 | 2098 | 0.175 | -0.284 | 0.044 | 1.11E-10 | ---- | 985 | -4.001 | 3.15E-05 |
|  |  | epi_high2 | 2141 | 0.172 | -0.275 | 0.044 | 3.57E-10 | ---- | 985 | -4.829 | 6.86E-07 |
|  |  | adp_high4 | 3147 | 0.139 | -0.241 | 0.040 | 1.16E-09 | ---- | 1858 | -4.761 | 9.63E-07 |
|  |  | epi_low4 | 2488 | 0.162 | -0.251 | 0.042 | 1.67E-09 | ---- | 1324 | -4.043 | 2.64E-05 |
|  |  | col_high2 | 3352 | 0.136 | 0.213 | 0.038 | 1.87E-08 | ++++ | 2002 | 4.223 | 1.21E-05 |
| 1:192194880 | G/C | epi_low5 | 3152 | 0.446 | 0.155 | 0.026 | 1.96E-09 | ++-+ | 1855 | 1.818 | 0.035 |
| 1:192195284 | T/C | epi_low5 | 3152 | 0.447 | 0.154 | 0.026 | 2.34E-09 | ++-+ | 1855 | 1.750 | 0.04 |
| 1:192182044 | A/C | epi_low5 | 3152 | 0.447 | 0.151 | 0.026 | 4.88E-09 | ++-+ | 1855 | 1.757 | 0.039 |
| 1:192193819 | T/C | epi_low5 | 3152 | 0.431 | 0.151 | 0.026 | 6.05E-09 | ++-+ | 1855 | 1.675 | 0.047 |
| 1:192206244 | C/T | epi_low5 | 3152 | 0.448 | 0.151 | 0.026 | 6.63E-09 | ++-+ | 1855 | 1.889 | 0.029 |
| 1:192168015 | T/A | epi_low5 | 3152 | 0.450 | 0.149 | 0.026 | 6.98E-09 | ++-+ | 1855 | 1.728 | 0.042 |
| 1:192142107 | T/TG | epi_low5 | 3152 | 0.469 | 0.145 | 0.026 | 1.45E-08 | +??+ | 1510 | 1.461 | 0.072 |
| 1:192148472 | C/T | epi_low5 | 3152 | 0.469 | 0.144 | 0.026 | 1.55E-08 | ++-+ | 1855 | 1.809 | 0.035 |
| 1:192154581 | A/G | epi_low5 | 3152 | 0.453 | 0.145 | 0.026 | 1.79E-08 | ++++ | 1855 | 1.798 | 0.036 |
| 1:192141301 | A/C | epi_low5 | 3152 | 0.436 | 0.146 | 0.026 | 1.81E-08 | ++-+ | 1855 | 1.666 | 0.048 |
| 1:192193726 | T/C | epi_low5 | 3152 | 0.413 | 0.147 | 0.026 | 1.84E-08 | ++-+ | 1855 | 2.154 | 0.016 |
| 1:192140500 | T/A | epi_low5 | 3152 | 0.437 | 0.145 | 0.026 | 2.17E-08 | ++-+ | 1855 | 1.667 | 0.048 |
| 1:192186870 | T/C | epi_low5 | 3152 | 0.412 | 0.146 | 0.026 | 2.37E-08 | ++-+ | 1855 | 2.144 | 0.016 |
| 1:192139135 | A/G | epi_low5 | 3152 | 0.461 | 0.143 | 0.026 | 2.51E-08 | ++-+ | 1855 | 1.693 | 0.045 |
| 1:192174246 | T/G | epi_low5 | 3152 | 0.417 | 0.145 | 0.026 | 2.76E-08 | ++-+ | 1855 | 2.120 | 0.017 |
| 1:192183215 | C/A | epi_low5 | 3152 | 0.412 | 0.144 | 0.026 | 3.67E-08 | ++-+ | 1855 | 2.155 | 0.016 |
| 1:192144114 | G/A | epi_low5 | 3152 | 0.442 | 0.141 | 0.026 | 4.57E-08 | ++-+ | 1855 | 2.132 | 0.017 |
| 1:192186916 | G/A | epi_low5 | 3152 | 0.411 | 0.143 | 0.026 | 4.61E-08 | ++-+ | 1855 | 2.164 | 0.015 |
| 5:19109993 | T/C | col_high1 | 3361 | 0.023 | 0.458 | 0.080 | 1.19E-08 | --?+ | 1909 | -2.090 | 0.982 |
|  |  | col_low2 | 3364 | 0.023 | 0.455 | 0.080 | 1.47E-08 | --?+ | 1909 | -1.567 | 0.941 |
| 6:121921871 | A/G | epi_low2 | 3027 | 0.085 | -0.273 | 0.049 | 2.22E-08 | ---+ | 1764 | 0.612 | 0.73 |
| 9:28873884 | T/A | epi_low2 | 3027 | 0.005 | -0.988 | 0.180 | 3.87E-08 | --?? | 1270 | -1.582 | 0.057 |
| 10:75490891 | A/G | col_high2 | 3352 | 0.006 | -0.858 | 0.152 | 1.52E-08 | -??+ | 1662 | 0.570 | 0.716 |
| 10:111139289 | T/A | epi_high3 | 3154 | 0.138 | -0.251 | 0.037 | 6.68E-12 | ---- | 1856 | -5.440 | 2.66E-08 |
|  |  | epi_low5 | 3152 | 0.137 | -0.249 | 0.037 | 9.70E-12 | ---- | 1855 | -5.124 | 1.50E-07 |
|  |  | epi_high2 | 2141 | 0.135 | -0.286 | 0.045 | 1.27E-10 | ---- | 985 | -3.844 | 6.05E-05 |
|  |  | epi_low4 | 2488 | 0.140 | -0.257 | 0.041 | 2.80E-10 | ---- | 1324 | -4.165 | 1.56E-05 |
|  |  | epi_high1 | 2098 | 0.137 | -0.280 | 0.045 | 3.43E-10 | ---- | 985 | -3.787 | 7.62E-05 |
|  |  | epi_low2 | 3027 | 0.137 | -0.235 | 0.037 | 3.44E-10 | ---- | 1764 | -4.816 | 7.32E-07 |
|  |  | epi_low3 | 2486 | 0.140 | -0.255 | 0.041 | 3.84E-10 | ---- | 1323 | -3.896 | 4.89E-05 |
|  |  | epi_low1 | 2962 | 0.134 | -0.210 | 0.038 | 3.70E-08 | ---- | 1689 | -4.452 | 4.25E-06 |
| 11:92185065 | A/T | adp_high2 | 2799 | 0.011 | -0.702 | 0.127 | 3.11E-08 | --?? | 1102 | -1.870 | 0.031 |
|  |  | adp_high1 | 2700 | 0.011 | -0.702 | 0.127 | 3.16E-08 | --?? | 1102 | -1.870 | 0.031 |
| 12:132589485 | G/A | col_high2 | 3352 | 0.010 | 0.669 | 0.121 | 2.88E-08 | ++?? | 1510 | 0.821 | 0.206 |
| 13:96912429 | A/G | epi_high1 | 2098 | 0.051 | -0.400 | 0.068 | 3.36E-09 | +++- | 985 | 0.653 | 0.743 |
| 17:16451482 | G/A | col_low2 | 3364 | 0.003 | 1.169 | 0.211 | 3.13E-08 | -??? | 1267 | -0.132 | 0.553 |
|  |  | col_high1 | 3361 | 0.003 | 1.155 | 0.211 | 4.39E-08 | -??? | 1267 | -0.132 | 0.553 |
| 17:21960955 | A/T | epi_low5 | 3152 | 0.274 | 0.260 | 0.029 | 7.73E-19 | (variant had imputation Rsq <0.7 in replication samples) | | | |
|  |  | epi_high3 | 3154 | 0.274 | 0.245 | 0.029 | 5.47E-17 |  |  |  |  |
|  |  | adp_low3 | 3014 | 0.271 | 0.167 | 0.030 | 2.87E-08 |  |  |  |  |
|  |  | adp_low1 | 3140 | 0.272 | 0.162 | 0.030 | 4.79E-08 |  |  |  |  |
| 18:29059923 | TAAATA/T | col_high1 | 3361 | 0.083 | -0.250 | 0.045 | 3.23E-08 | +??+ | 1662 | 0.549 | 0.292 |
|  |  | col_low2 | 3364 | 0.083 | -0.248 | 0.045 | 3.95E-08 | +??+ | 1662 | 0.759 | 0.224 |
| 20:50142397 | CTG/C | col_high2 | 3352 | 0.003 | 1.194 | 0.217 | 3.53E-08 | +??? | 1267 | 0.647 | 0.259 |

**Supplementary Table 3b** Discovery and Replication results of the variants that drove the gene-based signals. The variants were identified through leave-one-out analysis. Replication was performed as a meta-analysis in order of FHS, GeneSTAR European Americans, GeneSTAR African Americans and OOA for direction of effect. Replication p-value was computed using a one-sided test.

| **Gene** |  |  |  | **DISCOVERY** | | | | | **REPLICATION** | | | |
| --- | --- | --- | --- | --- | --- | --- | --- | --- | --- | --- | --- | --- |
|  | **snpID hg38** | **ref/alt** | **Phenotype** | **N** | **MAF** | **beta** | **se** | **pval** | **Direction** | **N** | **Zscore** | **pval** |
| *SVEP1* | 9:110549951 | G/C | adp_low1 | 3140 | 0.029 | 0.338 | 0.075 | 5.84E-06 | ++?+ | 1833 | 2.662 | 0.004 |
| *BCO1* | 16:81290348 | G/C | epi_low1 | 2962 | 0.007 | 0.719 | 0.147 | 1.05E-06 | --?? | 1195 | -0.575 | 0.717 |
| *IDH3A* | 15:78161708 | T/A | col_high2 | 3352 | 0.006 | 0.742 | 0.153 | 1.20E-06 | --?? | 1510 | -1.012 | 0.844 |

**Supplementary Table 4**: Platelet aggregation results in the Caerphilly Prospective Study.

Top, Lead variants and *RGS18* genome-wide significant variants. The italicized variant was the lead variant in the *RGS18* region discovery; Bottom, Variants driving gene-based SKAT.

|  | | | | | | **Trait: ADP (0.725uM)** | | | | **Trait: Collagen (42.7 ug/mL)** | | | |
| --- | --- | --- | --- | --- | --- | --- | --- | --- | --- | --- | --- | --- | --- |
| **snpID hg38** | **rsID** | **Ref/Alt** | **minor allele** | **MAF** | **Rsq** | **N** | **beta** | **se** | **pval** | **N** | **beta** | **se** | **pval** |
| 1:156899922 | rs12041331 | G/A | ALT | 0.080 | 0.992 | 1177 | -0.304 | 0.055 | 1.62E-08 | 811 | -0.688 | 0.145 | 9.75E-07 |
| 1:192194880 | rs1175170 | G/C | ALT | 0.487 | 0.987 | 1177 | 0.087 | 0.029 | 1.09E-03 | 811 | 0.203 | 0.075 | 3.52E-03 |
| 1:20567949 | rs12137738 | A/T | ALT | 0.108 | 0.961 | 1177 | -0.014 | 0.046 | 6.20E-01 | 811 | -0.020 | 0.126 | 5.64E-01 |
| 1:67128641 | rs142001088 | C/T | ALT | 0.023 | 0.887 | 1177 | -0.040 | 0.100 | 6.56E-01 | 811 | -0.500 | 0.259 | 9.73E-02 |
| 10:111139289 | rs7097060 | T/A | ALT | 0.188 | 0.988 | 1177 | -0.046 | 0.038 | 1.11E-01 | 811 | -0.074 | 0.100 | 2.31E-01 |
| 11:92185065 | rs183146849 | A/T | ALT | 0.019 | 0.863 | 1177 | 0.094 | 0.110 | 8.02E-01 | 811 | -0.444 | 0.297 | 9.33E-01 |
| 12:132589485 | rs140148392 | G/A | ALT | 0.014 | 0.934 | 1177 | 0.144 | 0.123 | 8.79E-01 | 811 | 0.506 | 0.322 | 9.42E-01 |
| 13:96912429 | rs61974290 | A/G | ALT | 0.067 | 0.963 | 1177 | 0.011 | 0.057 | 5.79E-01 | 811 | -0.054 | 0.151 | 6.40E-01 |
| 5:19109993 | rs112157462 | T/C | ALT | 0.028 | 0.892 | 1177 | 0.063 | 0.092 | 7.53E-01 | 811 | -0.259 | 0.260 | 1.59E-01 |
| 6:121921871 | rs58250884 | A/G | ALT | 0.052 | 0.997 | 1177 | -0.024 | 0.065 | 3.58E-01 | 811 | 0.126 | 0.180 | 7.59E-01 |
| 1:192139135 | rs12070423 | A/G | ALT | 0.496 | 0.983 | 1177 | 0.093 | 0.029 | 6.08E-04 | 811 | 0.239 | 0.076 | 8.29E-04 |
| 1:192140500 | rs10801100 | T/A | ALT | 0.496 | 0.985 | 1177 | 0.093 | 0.029 | 5.95E-04 | 811 | 0.239 | 0.076 | 7.93E-04 |
| 1:192141301 | rs7546592 | A/C | ALT | 0.496 | 0.985 | 1177 | 0.093 | 0.029 | 5.91E-04 | 811 | 0.239 | 0.076 | 7.96E-04 |
| 1:192144114 | rs10801102 | G/A | ALT | 0.479 | 0.986 | 1177 | 0.095 | 0.029 | 4.82E-04 | 811 | 0.221 | 0.076 | 1.81E-03 |
| 1:192148472 | rs6687273 | C/T | ALT | 0.495 | 0.994 | 1177 | 0.093 | 0.029 | 6.08E-04 | 811 | 0.233 | 0.075 | 9.91E-04 |
| 1:192154581 | rs7526348 | A/G | ALT | 0.495 | 0.998 | 1177 | 0.093 | 0.029 | 5.68E-04 | 811 | 0.235 | 0.075 | 9.03E-04 |
| 1:192168015 | rs10754003 | T/A | ALT | 0.493 | 0.993 | 1177 | 0.096 | 0.029 | 3.85E-04 | 811 | 0.235 | 0.075 | 8.86E-04 |
| 1:192174246 | rs12117018 | T/G | ALT | 0.480 | 0.992 | 1177 | 0.092 | 0.029 | 5.95E-04 | 811 | 0.205 | 0.075 | 3.17E-03 |
| 1:192182044 | rs1937235 | A/C | ALT | 0.487 | 0.989 | 1177 | 0.088 | 0.029 | 1.05E-03 | 811 | 0.203 | 0.075 | 3.54E-03 |
| 1:192183215 | rs10921107 | C/A | ALT | 0.473 | 0.989 | 1177 | 0.088 | 0.029 | 1.03E-03 | 811 | 0.183 | 0.075 | 7.42E-03 |
| 1:192186870 | rs2247567 | T/C | ALT | 0.473 | 0.989 | 1177 | 0.088 | 0.029 | 1.05E-03 | 811 | 0.183 | 0.075 | 7.42E-03 |
| 1:192186916 | rs2247566 | G/A | ALT | 0.471 | 0.988 | 1177 | 0.088 | 0.029 | 1.00E-03 | 811 | 0.180 | 0.076 | 8.67E-03 |
| 1:192193726 | rs1175168 | T/C | ALT | 0.472 | 0.988 | 1177 | 0.088 | 0.029 | 1.06E-03 | 811 | 0.184 | 0.075 | 7.34E-03 |
| 1:192193819 | rs1175169 | T/C | ALT | 0.487 | 0.988 | 1177 | 0.088 | 0.029 | 1.07E-03 | 811 | 0.204 | 0.075 | 3.40E-03 |
| *1:192194880* | *rs1175170* | *G/C* | *ALT* | *0.487* | *0.987* | *1177* | *0.087* | *0.029* | *1.09E-03* | *811* | *0.203* | *0.075* | *3.52E-03* |
| 1:192195284 | rs1175171 | T/C | ALT | 0.487 | 0.987 | 1177 | 0.087 | 0.029 | 1.09E-03 | 811 | 0.203 | 0.075 | 3.54E-03 |
| 1:192206244 | rs12037701 | C/T | REF | 0.490 | 0.982 | 1177 | -0.087 | 0.028 | 1.08E-03 | 811 | -0.150 | 0.075 | 2.25E-02 |

| **Gene** | **snpID hg38** | **rsID** | **REF** | **ALT** | **minor allele** | **MAF** | **Rsq** | **Trait** | **N** | **beta** | **Se** | **pval** |
| --- | --- | --- | --- | --- | --- | --- | --- | --- | --- | --- | --- | --- |
| SVEP1 | 9:110549951 | rs61751937 | G | C | ALT | 0.018 | 0.999 | ADP | 1177 | 0.251 | 0.104 | 7.98E-03 |
| BCO1 | 16:81290348 | rs143238312 | G | C | ALT | 0.013 | 0.989 | ADP | 1177 | 0.245 | 0.128 | 2.74E-02 |
| BCO1 | 16:81290348 | rs143238312 | G | C | ALT | 0.013 | 0.989 | Thrombin | 1183 | 0.365 | 0.163 | 1.27E-02 |

**Supplementary Table 5:** Co-localization between all transcripts having a platelet eQTL p-value <0.0031(0.05/16) within +/- 20KB of GWAS locus peak. Meaningful co-localization was noted between PEAR1 and RGS18 for Chr1:156899922 and Chr1:192194880 loci, respectively.

| **Locus** | **Phenotype** | **Co-localization by gene (posterior probability in %)** |
| --- | --- | --- |
| Chr1:20567949 | epi_low3 | NBPF3 (9.54%) |
| Chr1:67128641 | col_high1 | SLC35D1 (0.629%) |
| Chr1:156899922 | adp_low1 | LMNA (2.64%), **PEAR1 (99.6%),** |
| Chr1:156899922 | col_high1 | LMNA (2.64%), **PEAR1 (99.6%)** |
| Chr1:156899922 | epi_low1 | LMNA (2.64%), **PEAR1 (99.6%)** |
| Chr1:192194880 | adp_high1 | **RGS18 (69.7%)** |
| Chr1:192194880 | epi_low5 | **RGS18 (69.0%)** |
| Chr5:19109993 | col_high1 | - |
| Chr6:121921871 | epi_low2 | - |
| Chr9:28873884 | epi_low2 | - |
| Chr10:75490891 | col_high2 | - |
| Chr10:111139289 | epi_high3 | ADRA2A (34.5%) |
| Chr11:92185065 | adp_high2 | - |
| Chr12:132589485 | col_high2 | ANKLE2 (0.0818%), FBRSL1 (0.115%) |
| Chr13:96912429 | epi_high1 | - |
| Chr17:21960955 | epi_low5 | - |
| Chr17:21960955 | adp_low3 | - |
| Chr17:16451482 | col_low2 | LLRC75A-AS1 (1.67%) |
| Chr18:29059923 | col_low2 | - |
| Chr20:50142397 | col_high2 | BCAS4 (0.369%) |

**Supplementary Table 6:** UKBB was queried for the top 16 independent variants. Only blood and cardiovascular traits are shown here. NA: the variant is not available in UKBB for lookup.

| **Locus** | **rsID** | **Effect allele** | **UKBB (p-value, beta)** |
| --- | --- | --- | --- |
| Chr1:192194880 | rs1175170 | C | Mean platelet volume (0.00012, 0.006), Platelet distribution width (0.0039, 0.002), Platelet count (0.0044, -0.262), Cerebral infarction (0.0067, 0.0004), Monocyte count (0.0083, -0.0007), Basophil percentage (0.015, -0.002), Subarachnoid hemorrhage (0.025, -0.0002) |
| Chr1:156899922 | rs12041331 | A | Mean platelet volume (2.46E-132, -0.070), Platelet distribution width (4.16e-31, -0.018), Platelet count (1.90E-27, 1.793), I20-I25 Ischemic heart diseases (0.0016, -0.002), Stroke (0.0028, -0.0005), heart/cardiac problem (0.0041, -0.0026), angina (0.0049, -0.001), Chronic ischemic heart disease (0.0090, -0.002), heart attack/myocardial infarction (0.012, -0.001), Reticulocyte percentage (0.016, -0.004), Reticulocyte count (0.016, -0.0001), Monocyte percentage (0.023, 0.015), Hematocrit percentage (0.024, 0.020) |
| Chr1:20567949 | rs12137738 | T | Eosinophil percentage (0.013, -0.013), Mean corpuscular hemoglobin (0.017, -0.011), Mean reticulocyte volume (0.024, -0.050) |
| Chr1:67128641 | rs142001088 | T | NA |
| Chr5:19109993 | rs112157462 | C | Retinal vascular occlusions (0.019, 0.0005), other venous/lymphatic disease (0.022, 0.0009), varicose veins (0.023, 0.0008), White blood cell count (0.035, -0.020), Platelet crit (0.037, -0.0004), pulmonary embolism +/- dvt (0.051, 0.001) |
| Chr6:121921871 | rs58250884 | G | Lymphocyte count (0.0033428, 0.007), White blood cell count (0.025, 0.016) |
| Chr9:28873884 | rs185159562 | A | NA |
| Chr10:75490891 | rs138028657 | G | NA |
| Chr10:111139289 | rs7097060 | A | Occlusion and stenosis of prevertebral arteries (0.0025, 0.0004), Lymphocyte percentage (0.0035, 0.052), Neutrophil percentage (0.016, -0.049), Nucleated red blood cell count (0.047, 0.0001), Lymphocyte count (0.048, 0.002) |
| Chr11:92185065 | rs183146849 | T | Other forms of heart disease (0.014, 0.005) |
| Chr12:132589485 | rs140148392 | A | NA |
| Chr13:96912429 | rs61974290 | G | Lymphocyte count (0.0037, -0.006), G Red blood cell distribution width (0.0094, -0.008), Stroke (0.012, 0.0005), White blood cell count (0.012, -0.014), other venous/lymphatic disease (0.012, 0.0006), Mean reticulocyte volume (0.016, 0.059), Varicose veins of lower extremity (0.017, -0.001), Monocyte count (0.020, -0.001), Platelet count (0.045, -0.343) |
| Chr17:21960955 | . | T | NA |
| Chr17:16451482 | rs575524466 | A | NA |
| Chr18:29059923 | rs138845468 | T | NA |
| Chr:2050142397 | rs542707094 | C | NA |

**Supplementary Table 7**: Annotation of genome-wide significant variants in RGS18 locus. Annotation evidence for megakaryocyte (MK) elements arbitrarily labeled to indicate four independent enhancer regions (e11 – e14) falling within a common super enhancer (se11).

|  | | **Lowest P value for Trait** | | | **Encode Regulatory** | | **MK specific Regulatory** | | **Platelet RGS18 eQTL** | | |
| --- | --- | --- | --- | --- | --- | --- | --- | --- | --- | --- | --- |
| **snpID hg38** | **MAF** | **ADP** | **COL** | **EPI** | **DNAse** | **TFBS Cluster** | **Enhancer** | **Super Enhancer** | **Beta** | **Var** | **Pval** |
| 1:192139135:A:G | 0.456 | 1.16E-05 | 4.42E-02 | 2.51E-08 | Yes | GATA1 | . | . | 0.06 | 0.00 | 3.13E-02 |
| 1:192140500:T:A | 0.430 | 3.54E-06 | 2.54E-02 | 2.17E-08 | Yes | . | . | . | 0.06 | 0.00 | 3.13E-02 |
| 1:192141301:A:C | 0.430 | 2.55E-06 | 2.51E-02 | 1.81E-08 | No | . | . | . | 0.06 | 0.00 | 3.13E-02 |
| 1:192142107:T:TG | 0.466 | 1.56E-05 | 5.81E-02 | 1.45E-08 | No | . | . | . | . | . | . |
| 1:192144114:G:A | 0.437 | 3.34E-06 | 8.53E-02 | 4.57E-08 | No | . | . | . | 0.06 | 0.00 | 2.71E-02 |
| 1:192148472:C:T | 0.465 | 1.17E-05 | 4.08E-02 | 1.55E-08 | Yes | FOS\|CEBPB | e11 | se11 | 0.06 | 0.00 | 3.13E-02 |
| 1:192154581:A:G | 0.449 | 2.79E-06 | 3.57E-02 | 1.79E-08 | Yes | . | e12 | se11 | 0.06 | 0.00 | 3.18E-02 |
| 1:192168015:T:A | 0.445 | 9.49E-06 | 4.20E-02 | 6.98E-09 | Yes | CTCF | e13 | se11 | 0.06 | 0.00 | 2.73E-02 |
| 1:192174246:T:G | 0.410 | 5.36E-07 | 5.55E-02 | 2.76E-08 | No | . | e14 | se11 | 0.07 | 0.00 | 1.40E-02 |
| 1:192182044:A:C | 0.443 | 1.81E-05 | 3.71E-02 | 4.88E-09 | No | . | . | . | 0.08 | 0.00 | 2.29E-03 |
| 1:192183215:C:A | 0.406 | 1.14E-06 | 5.63E-02 | 3.67E-08 | No | . | . | . | 0.08 | 0.00 | 1.83E-03 |
| 1:192186870:T:C | 0.405 | 2.02E-06 | 5.64E-02 | 2.37E-08 | Yes | . | . | . | 0.08 | 0.00 | 1.83E-03 |
| 1:192186916:G:A | 0.405 | 9.19E-07 | 6.08E-02 | 4.61E-08 | Yes | . | . | . | 0.09 | 0.00 | 1.01E-03 |
| 1:192193726:T:C | 0.407 | 1.11E-06 | 5.35E-02 | 1.84E-08 | No | . | . | . | 0.08 | 0.00 | 1.83E-03 |
| 1:192193819:T:C | 0.424 | 3.79E-06 | 3.76E-02 | 6.05E-09 | No | . | . | . | 0.08 | 0.00 | 2.29E-03 |
| 1:192194880:G:C | 0.442 | 7.86E-06 | 2.37E-02 | 1.96E-09 | Yes | . | . | . | 0.08 | 0.00 | 2.29E-03 |
| 1:192195284:T:C | 0.443 | 1.23E-05 | 3.47E-02 | 2.34E-09 | No | . | . | . | 0.08 | 0.00 | 2.38E-03 |
| 1:192206244:C:T | 0.441 | 5.94E-07 | 1.32E-02 | 6.63E-09 | Yes | E2F1 | . | . | 0.06 | 0.00 | 9.15E-03 |

**Supplementary Table 8:** Aggregated rare deleterious coding variants of 4 genes (SVEP1, BCO1, NELFA and IDH3A) were associated with platelet aggregation after Bonferroni correction (0.05 / 17744 = 2.819E-6) by SKAT with MAF threshold 0.05.

| **Gene** | **Chr** | **Start** | **Stop** | **# variants** | **p-value** | **Trait** |
| --- | --- | --- | --- | --- | --- | --- |
| *SVEP1* | 9 | 110365251 | 110579880 | 64 | 2.64E-06 | adp_low1 |
| *BCO1* | 16 | 81238448 | 81291142 | 27 | 8.88E-07 | epi_low1 |
| *NELFA* | 4 | 1982714 | 2041903 | 11 | 1.70E-06 | col_high1 |
| *IDH3A* | 15 | 78131498 | 78171949 | 10 | 2.40E-06 | col_high2 |

**Supplementary Table 9:** Annotation of 5 variants driving signals of SVEP1, BCO1, NELFA and IDH3A identified by SKAT gene-based test with MAF threshold 0.05.

| **Chr:pos** | **Ref/Alt** | **Gene** | **AA Change** | **dbSNP** | **SIFT** | **P2 HVAR** | **LRT** | **Mutation Taster** | **Meta**  **SVM** | **M-CAP** | **CADD** | **REVEL** |
| --- | --- | --- | --- | --- | --- | --- | --- | --- | --- | --- | --- | --- |
| 16:81290348 | G/C | BCO1 | p.G472A | rs143238312 | D | D | D | D | D | . | 25.2 | 0.875 |
| 9:110549951 | G/C | SVEP1 | p.R229G | rs61751937 | D | D | D | D | D | . | 28.3 | 0.672 |
| 4:1986122 | T/C | NELFA | p.K287R | rs150291014 | T | B | N | D | T | D | 15.89 | 0.062 |
| 4:1987948 | G/A | NELFA | p.R213W | rs763817905 | D | D | D | D | D | D | 35 | 0.457 |
| 15:78161708 | T/A | IDH3A | p.D139E | rs61752770 | D | P | D | D | T | D | 24.7 | 0.208 |

**Supplementary Table 10:** UKBB was queried for rs111245230 and rs61751937 found within *SVEP1*.

| **rsID** | **Effect allele** | **UKBB (p-value, beta)** |
| --- | --- | --- |
| rs61751937 | C | Mean platelet volume (0.00012, 0.006), Platelet distribution width (0.0039, 0.002), Platelet count (0.0044, -0.262), Cerebral infarction (0.0067, 0.0004), Monocyte count (0.0083, -0.0007), Basophil percentage (0.015, -0.002), Subarachnoid hemorrhage (0.025, -0.0002) |
| rs111245230 | T | Hypertension(1.14E-15, -0.017), White blood cell count(1.27E-05, -0.036), Neutrophil count(0.00035, -0.024), Platelet count(0.00056, -0.857),Monocyte count(0.0014, -0.002), high cholesterol(0.0026, -0.005), Mean reticulocyte volume(0.0045, 0.100), I21 Acute myocardial infarction(0.0057, -0.002), Eosinophil count(0.0062, -0.001), Platelet crit(0.0075, -0.0005), Body fat percentage(0.011, -0.074) , Ischemic heart diseases(0.018, -0.003), Intracerebral hemorrhage(0.021, -0.0005), Chronic ischemic heart disease (0.024, -0.002), Lymphocyte count(0.033, -0.006) |
